# Transcriptomics Unveil Canonical and Non-Canonical Heat Shock-Induced Pathways in Human Cell Lines

**DOI:** 10.1101/2024.12.22.629972

**Authors:** Andrew Reinschmidt, Luis Solano, Yonny Chavez, William Drew Hulsy, Nikolas Nikolaidis

## Abstract

The cellular stress response (CSR) is a conserved mechanism that protects cells from environmental and physiological stressors. The heat shock response (HSR), a critical component of the CSR, utilizes molecular chaperones to mitigate proteotoxic stress caused by elevated temperatures. We hypothesized that while the canonical HSR pathways are conserved across cell types, specific cell lines may exhibit unique transcriptional responses to heat shock. To test this, we compared the transcriptomic responses of HEK293, HepG2, and HeLa cells under control conditions immediately following heat shock and after an 8-hour recovery period. RNA sequencing revealed conserved activation of canonical HSR pathways, including the unfolded protein response, alongside enrichment of the non-canonical “Receptor Ligand Activity” pathway across all cell lines. Cell line-specific variations were also observed, with HepG2 cells displaying more uniquely expressed genes and elevated expression levels (fold changes) of shared genes under stress conditions. Validation by qPCR confirmed the activation of key genes within the “Receptor Ligand Activity” pathway across time points. These findings provide insights into conserved and context-specific aspects of the HSR, contributing to a more comprehensive understanding of stress response mechanisms across mammalian cells.

## Introduction

The cellular stress response (CSR) is a critical survival mechanism that helps cells cope with environmental stressors, and the heat shock response (HSR) is one of its most essential components [1-7]. The HSR is primarily activated by temperature increases and other stressors, triggering the activation of heat shock factors (HSFs), especially HSF1, which then drive the expression of heat shock proteins (HSPs). These molecular chaperones, including HSP70 and HSP90, facilitate protein homeostasis by refolding misfolded proteins, preventing aggregation, and modulating apoptosis [3-5,8-12]. While the canonical HSR is well-characterized, the full scope of its regulatory mechanisms and potential adaptations to different cellular contexts still need to be understood [13]. In particular, non-canonical HSR pathways and their roles in cellular stress adaptation are less explored [14], despite their potential significance in various disease processes.

In cancer, the HSR’s molecular machinery is hijacked for cellular survival, enabling tumor cells to withstand the proteotoxic stress generated by rapid proliferation, hypoxia, and therapeutic interventions. Cancer cells frequently upregulate key HSR components, including HSF1, to maintain proteostasis and resist apoptosis induced by therapies [15-18]. While much is known about the activation of canonical HSR pathways in cancer, there is a significant gap in understanding how these pathways are modulated in cancer cells and whether non-canonical HSR pathways contribute to the enhanced stress resistance observed in tumors. Current research broadly focuses on the well-established roles of HSPs and HSF1. Still, the complexity of the HSR—particularly its dynamic crosstalk with other stress response networks like the unfolded protein response (UPR) and DNA damage response (DDR)—and how these interactions regulate stress resistance in tumors remain underexplored [1-12].

This study aims to fill these gaps by conducting a comprehensive RNA sequencing (RNA-seq) analysis comparing cancerous (HeLa, HepG2) and non-cancerous (HEK293) cell lines exposed to heat stress. By comparing these cell lines, we aim to identify both conserved and cell-specific mechanisms regulating the HSR, with particular emphasis on uncovering non-canonical pathways that may play a significant role in cellular stress adaptation. We hypothesize that, while canonical HSR pathways are activated across all cell types, cancer cells will exhibit unique regulatory modifications and interactions, particularly involving non-canonical HSR pathways. These candidate pathways may contribute to enhanced survival and therapy resistance phenotypes. Specifically, we expect to observe differential regulation of key HSR components, novel interactions between the HSR and other stress response pathways (e.g., UPR, DDR), and the activation of non-canonical HSR targets critical for cellular stress adaptation in cancer cells.

By identifying these novel non-canonical HSR targets, we anticipate uncovering new insights into how cells adapt to stress, particularly in the context of cancer. Moreover, our study will provide a deeper understanding of the full spectrum of HSR mechanisms, including those understudied. This research has broad implications not only for cancer biology but also for understanding the role of stress responses in a variety of other diseases, including neurodegenerative disorders and aging, where similar proteotoxic stress is a hallmark [19,20]. By uncovering the molecular mechanisms driving non-canonical HSR pathways, this work can potentially inform the development of novel therapeutic strategies that target stress responses, providing new avenues for treating cancer and other stress-related diseases.

## Materials and Methods

### Cell Culture

To examine the differential heat shock response, we utilized three human cell lines: human embryonic kidney cells (HEK293; ATCC® CRL-1573™), HeLa cells derived from Henrietta Lacks (ATCC® CCL-2™), and hepatocellular carcinoma cells (HepG2; ATCC® HB-8065) obtained from ATCC in December 2016 and verified bi-annually. HEK293 cells were cultured in Dulbecco’s Modified Eagle Medium (DMEM), while HeLa and HepG2 cells were grown in Minimum Essential Medium (MEM). Media for both cell lines were supplemented with 10% fetal bovine serum (FBS), 2 mM L-glutamine, and penicillin-streptomycin. HeLa and HepG2 media were supplemented with 0.1 mM non-essential amino acids (NEAA) and 1 mM sodium pyruvate. All cultures were maintained in a humidified atmosphere containing 5% CO2 at 37°C.

### Heat Shock Treatment

To assess the effect of heat shock on transcription, cells were either maintained at 37°C or subjected to heat stress at 42°C for 60 minutes in a humidified CO2 incubator. Following heat shock, cells were allowed to recover at 37°C for 8 hours. This procedure was independently conducted on HeLa, HEK293, and HepG2 cell lines. Newly thawed, low-passage cells (passages 4–7) were cultured in 175 cm^2^ flasks until reaching approximately 80% confluency. For each experimental condition, three flasks were used: one remained at 37°C as a control, while two others were exposed to 42°C for one hour to induce heat shock. Of the heat-shocked flasks, one was processed immediately after heat shock (0 hours recovery, 0R), and the other was allowed to recover at 37°C for 8 hours (8 hours recovery, 8R) (Figure 1). This procedure was repeated using freshly thawed cells to generate biological replicates. After treatment, cells were harvested through trypsinization, pelleted, and frozen for subsequent analysis.

**Figure 1.**
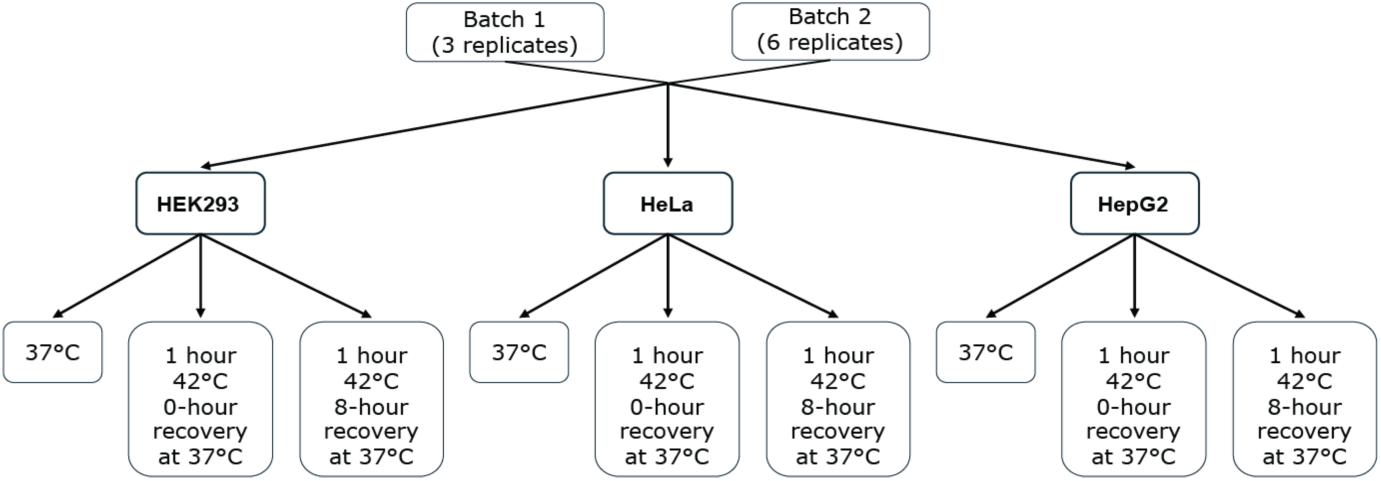
Flowchart diagram illustrating the experimental design and treatment conditions. Two experimental batches were analyzed, consisting of 3 and 6 replicates for HEK293, HeLa, and HepG2 cell lines. Samples were subjected to a 1-hour mild heat shock at 42°C and allowed to recover for either 0 hours or 8 hours at 37°C. Control samples were maintained at 37°C for the entire experiment. This experimental design thoroughly compares gene expression changes induced by heat shock across replicates and batches.

### Sample Preparation, cDNA Library Preparation, and Sequencing

Cells from the control, 0R, and 8R conditions were sent to Novogene (Sacramento, CA) for RNA isolation, library preparation, and sequencing. Total RNA was isolated using the Qiagen RNeasy series kit, and quality control (QC) was meticulously performed at each step to ensure reliable data. RNA degradation and contamination were assessed on 1% agarose gels, purity was verified using a NanoPhotometer® spectrophotometer (IMPLEN, CA, USA), and integrity and quantification were evaluated using the RNA Nano 6000 Assay Kit on the Bioanalyzer 2100 system (Agilent Technologies, CA, USA). For transcriptome sequencing, 1 μg of total RNA per sample was used to generate sequencing libraries with the NEBNext® UltraTM RNA Library Prep Kit for Illumina® (NEB, USA) following the manufacturer’s protocol. mRNA was isolated using poly-T oligo-attached magnetic beads, fragmented under elevated temperature, and reverse transcribed into cDNA. First and second-strand cDNA synthesis involved random hexamer primers, M-MuLV Reverse Transcriptase, DNA Polymerase I, and RNase H, followed by end repair, adenylation, and ligation of NEBNext adaptors. cDNA fragments of 150–200 bp were selected using AMPure XP beads (Beckman Coulter, USA), processed with USER Enzyme, and amplified via PCR with Phusion High-Fidelity DNA polymerase and indexed primers. Libraries were purified with the AMPure XP system and validated on the Agilent Bioanalyzer 2100 system. Clustering was conducted on a cBot Cluster Generation System using a PE Cluster Kit cBot-HS (Illumina), and paired-end sequencing was performed on an Illumina NovaSeq S4 PE100 platform, generating at least 30 million reads per sample across all conditions.

### Transcriptomics Analyses (detailed methodology in Appendix 1)

#### Read Quality Control Overview

Adapter sequences, PhiX library sequences, and low-quality reads are common byproducts of library preparation and next-generation sequencing (NGS). To ensure that high-quality reads are used for downstream analysis, BBDuk was employed to perform adapter trimming, PhiX filtering, and quality filtering based on Phred scores [21,22].

#### Adapter and PhiX Trimming

To eliminate unwanted sequences from downstream analysis, we used BBDuk’s k-mer trimming functionality for both adapter and PhiX filtering. Each paired read was processed with a BBDuk shell script to remove reads that matched reference k-mers associated with adapter sequences or the PhiX control library.

#### Phred Quality Score

To ensure high-quality reads were retained, BBDuk’s Phred quality trimming function was applied. This function scans the right end of paired reads, trimming until the Phred quality score meets the specified threshold. If the score remains unsatisfactory, the entire paired read is discarded. These processes were executed via command line for each pair of reads.

#### Reference Genome Indexing and Alignment of Reads to an Indexed Reference Genome

To quantify gene expression, it is essential to map high-quality reads to the human reference genome to identify the features in the dataset. STAR was employed for this purpose, accepting reads cleaned by BBDuk and outputting binary sequence alignment map files (.BAM) after performing splice-aware mapping against the indexed human reference genome (GRCh38.p13) [23]. Before alignment, the indexed reference genome was generated.

#### Sort and Index Mapped Reads for Feature Counting

Samtools was used to sort an input.BAM file and create a corresponding index file (.BAI), which is necessary for downstream feature counting steps [24]. The splice-aware and mapped.BAM files generated by STAR, as detailed in “Reference Genome Indexing and Alignment of Reads to an Indexed Reference Genome,” served as the input.

#### Feature Counts

HTSeq was utilized to create a raw gene count matrix by determining the number of reads mapped to each feature. The inputs required for HTSeq include a sorted aligned.BAM file, its corresponding index.BAI file, and a reference human genome annotation file (GRCh38.104.gtf) [25]. The sorted and indexed alignment files were produced as outlined in “Sort and Index Mapped Reads for Feature Counting.”

#### Quality Control

BBDuk, STAR, Samtools, and HTSeq generated multiple output files distributed across various directories. To summarize and evaluate these files, MultiQC was utilized, which provides an overview of file types associated with next-generation sequencing read processing [26]. By running MultiQC from a directory containing all relevant subdirectories with output files, users can verify the expected number of files and their types for each sample. Supplementary Tables 1 and 2 display the quality control metrics for each sample in batch 1 and batch 2, respectively.

#### Generating and Visualizing Differentially Expressed Genes

The negative binomial distribution is a suitable model for analyzing raw count data that is well-dispersed and influenced by multiple sources of variance, making DESeq2 an ideal choice for identifying differentially expressed genes (DEGs) [27]. DESeq2 estimates the mean and models the dispersion of gene expression for each gene observed in the dataset.

#### Gene Counts Normalization and Dimensionality Reduction

Utilizing DESeq2 for differential gene expression (DGE) analysis allows researchers to leverage its built-in variance stabilizing transformation (VST), which helps reduce the impact of outliers and addresses heteroscedasticity [27]. The VST-normalized counts were subsequently used as inputs for dimensionality reduction analyses. Principal component analysis (PCA) and row-scaled heatmap analysis were employed as dimensionality reduction techniques. These methods were chosen to effectively visualize the intricate VST-normalized gene expression data, encompassing over 20,000 gene features across nine samples, into a more concise and interpretable format. PCA was conducted to identify the primary sources of variation within the dataset and to estimate the effect sizes between samples based on the gene features contributing to that variance. PCA was conducted using R’s prcomp function, along with the tidyverse and ggfortify libraries.

#### Histograms, Volcano Plots, and Venn Diagrams

Differentially expressed genes were acquired from the normalized log2FC data created by DESeq2. Separating by cell line, the over and under-expressed genes for each condition comparison (0RvsControl, 8RvsControl, 8Rvs0R) were categorized as log2FC > 0.5 and log2FC < 0.5, respectively. These were plotted on a histogram in R using ggplot2 to visualize the number of DEGs present within a cell line at each pairwise condition comparison. Volcano plots were then generated for each condition pairwise comparison with R’s ggplot2 library to visualize statistically significant fold changes in gene expression. To visualize the significant batch conserved DEGs within each cell line at a given condition, Venn diagrams were created using tidy verse’s ggplot2 library. Significant DEGs were defined as having a |log2FC| > 0.5 and a p.adj.<0.05 (Benjamini-Hochberg). Tables with DEGs from all comparisons are in Appendix 2.

#### Functional Enrichment Analysis Using GSEA

Gene Set Enrichment Analysis (GSEA) was run against the ontological gene set collections (C5) defined by the molecular signatures database [28,29]. GSEA inputs ranked log2FC outputs from DESeq2 and designates an enrichment score for each pathway within the gene set collection. Enrichment scores are defined by increases to a running-sum statistic when a gene is in the gene set and decreases when it is not; the normalized enrichment scores (NES) then enable comparison across gene sets by accounting for differences in set size and correlations between the gene set and expression dataset. For this study, only human collections were used. For this reason, the normalized enrichment score was used in this analysis.

#### Functional Enrichment Analysis using STRING

STRING analysis investigated protein-protein interactions across various batches, cell lines, and condition comparisons. The STRING database aggregates, scores, and integrates publicly available information on protein-protein interactions, leveraging these data sources to generate predictive models of interaction networks [30]. We used our differentially expressed gene (DEG) data obtained from DESeq2 to carry out an enrichment analysis. To determine conserved pathways, the tables were compared, and genesets present in both batches and all three cell lines at a given condition were compiled in a table.

#### Visualization of Gene Expression Patterns Using Heatmaps

Row scaled heatmaps built with the Complexheatmap library were used to visualize the VST normalized expression patterns of genes found in “Heat Acclimation” and “Receptor Ligand Activity” gene sets. Thirteen genes from “Receptor Ligand Activity” with conserved gene expression in both batches, three cell lines, and three condition pairwise comparisons were identified. All six genes from “Heat Acclimation” were visualized with row scaled heatmaps for each batch [27,31].

#### Cytoscape Generated Network

Cytoscape’s [32] predicted network analysis was performed to determine the interactions of genes based on the acquired expression data. The network analysis uses a list of DEGs and the known interaction networks in the MSigdbr database.

#### Molecular Validation of using qPCR

Following the manufacturer’s protocol, RNA was isolated using 4 million HeLa cells per condition (control cells, 0-hour recovery, 8-hour recovery; different batches from the ones used for RNA-seq) using the Direct-Zol RNA mini-prep Kit (ZymoResearch, Irvine, CA, USA). Following the manufacturer’s protocol, cDNA was synthesized from 1ug of total RNA using the Superscript IV First-Strand synthesis system (ThermoFisher Scientific) and Oligo(dT)_20_ primers. cDNA samples were diluted to a concentration of 50 ng/uL. qPCR reactions were prepared with the Power SYBR™ Green PCR Master Mix (ThermoFisher Scientific, Waltham, MA, USA) according to the manufacturer’s instructions. Three biological replicates were run for each gene and condition [Gene names and primers (generated using NCBI’s primer-blast utility) are shown in Supplementary Table 3]. qPCR was performed using the CFX96 Touch Real-Time Detection System (Bio-Rad, Hercules, CA, United States). The relative normalized expression [33] of the raw transcript levels was calculated using the Livak method for each gene [34] using the software provided with the instrument [35]. The reference genes used in this method were ACTB and GAPDH. Statistical significance was assessed using one-way ANOVA (Analysis of Variance) followed by post-hoc Tukey HSD (Honestly Significant Difference) and Bonferroni tests. A P value < 0.05 was considered statistically significant. Results were plotted via boxplot using BoxPlotR [36].

## Results

### Principal Component Analysis

To identify key drivers of variance in the transcriptomic dataset, we conducted a principal component analysis (PCA) using VST-normalized read counts from three cell lines (HEK293, HepG2, and HeLa) under three conditions (Control, 0 hours recovery after heat shock, and 8 hours recovery after heat shock) across two experimental batches (Batch 1 and Batch 2). PCA results reveal that samples primarily cluster by cell line, regardless of heat shock condition or batch, indicating that cell line type is the most significant source of variance in gene expression (Figure 2).

**Figure 2.**
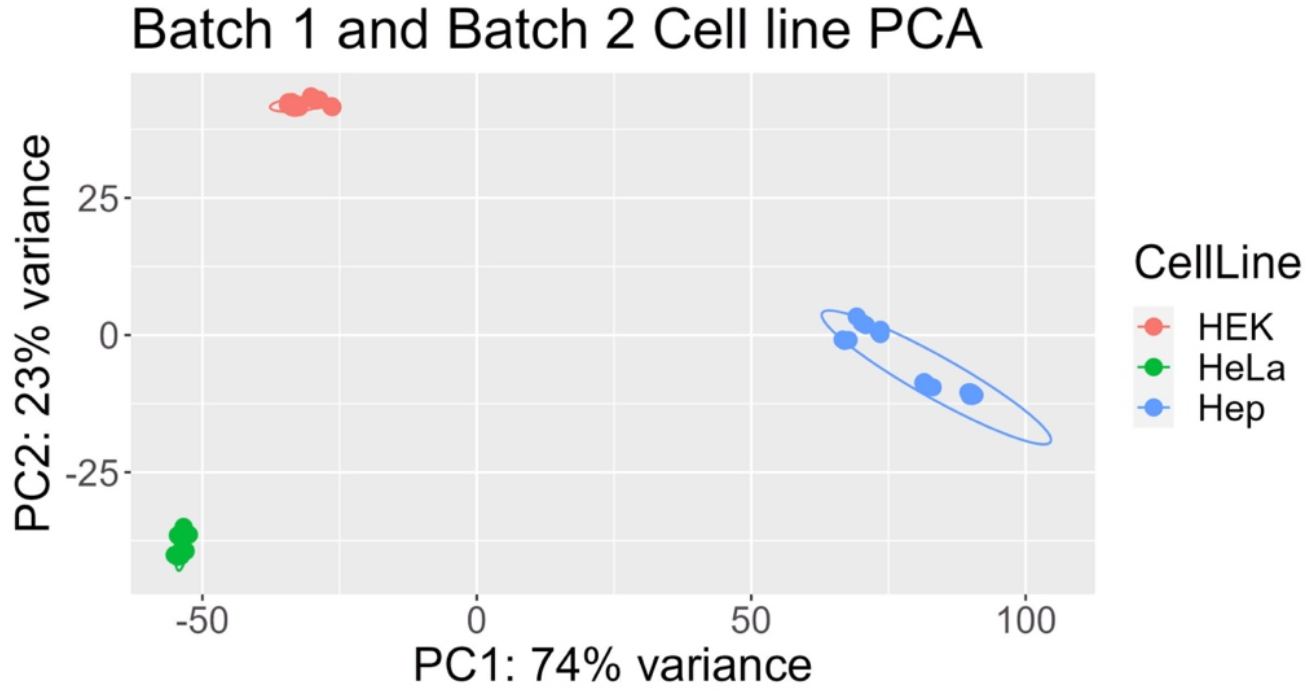
Principal component analysis (PCA) of variance-stabilizing transformed (VST) counts reveals that the primary source of variance in the dataset is the cell line type, regardless of batch or heat shock treatment. PCA reduces high-dimensional gene expression data to principal components, visually representing clustering patterns based on sample similarities. Each point represents a sample, with closer proximity indicating higher similarity.

Further PCA analyses on individual cell lines show that within-cell line variance is predominantly driven by experimental batch effects (Supplementary Figure 1). These findings necessitated batch-specific analyses to mitigate confounding effects.

When analyzing data separated by cell line and batch, PCA revealed that heat shock conditions significantly contributed to gene expression variance (Figure 3). For the HEK293 cell line, heat shock (0h recovery) explained 81% of the variance in Batch 1 (and 65% in Batch 2; Figure 3A, D), emphasizing a robust transcriptional response immediately following stress. For HeLa cells, heat shock accounted for 77% of the variance for Batch 1, while recovery from heat shock could explain 53% of the variance for Batch 2 (Figure 3B, E). HepG2 cells displayed a distinct pattern: most of the variance was explained by recovery from heat shock (8h), with PC1 capturing 80% of the variance in Batch 1 (and 78% in Batch 2; Figure 3C, F). The observed pattern highlights cell-line-specific differences in the heat shock response (HSR). These findings collectively establish that cell type, batch, and heat shock are the principal drivers of transcriptomic variance while revealing unique stress response and recovery dynamics across cell lines.

**Figure 3.**
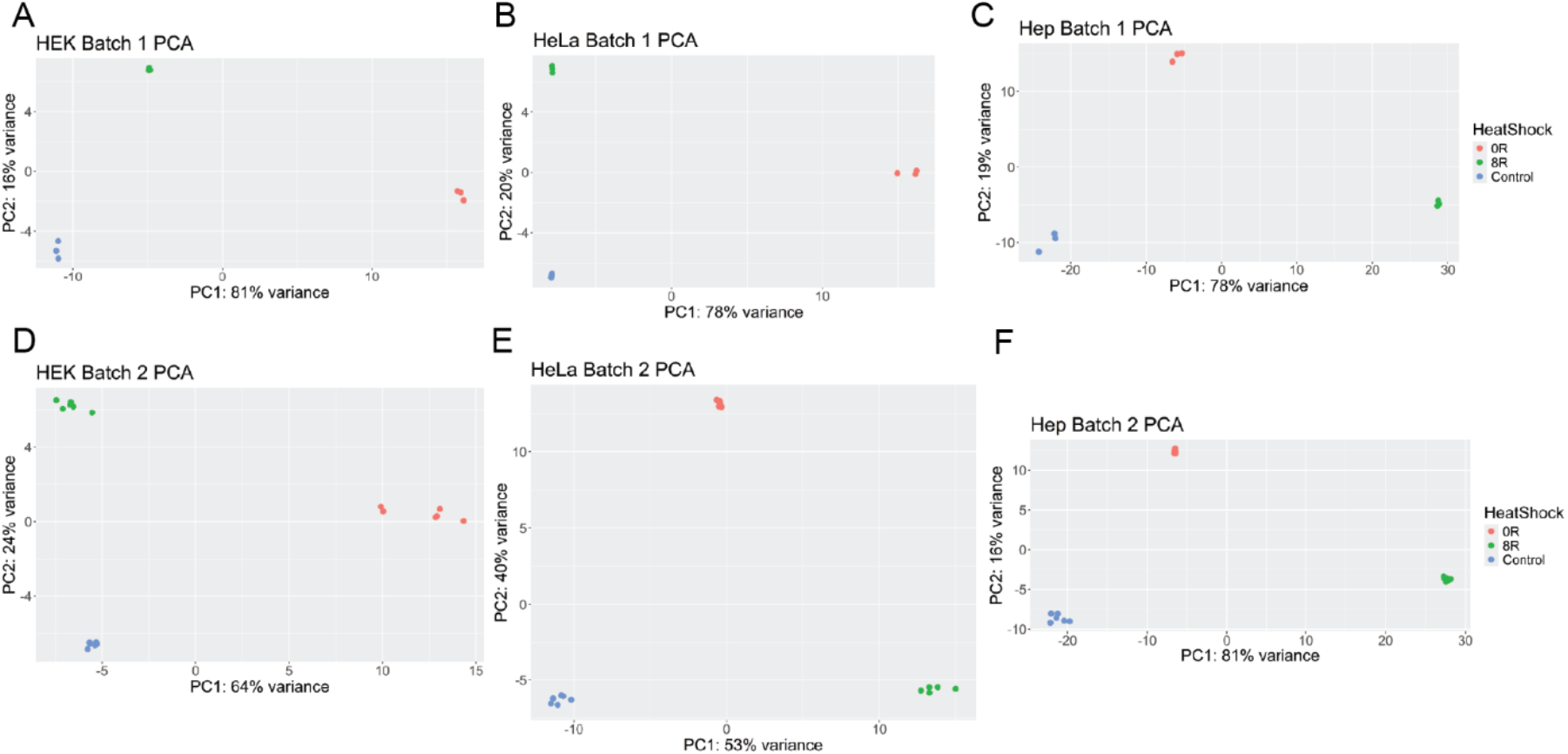
PCA of VST-normalized counts, stratified by batch and cell line, highlights clustering based on heat shock conditions. Individual PCA plots for HEK293 (Figures 3A, 3D), HeLa (Figures 3B, 3E), and HepG2 (Figures 3C, 3F) show distinct groupings between heat-shocked and control samples, indicating a robust transcriptional response to heat shock across all cell lines. The separation along principal components reflects the contribution of gene expression differences induced by heat shock and recovery time.

### Differential Gene Expression Analysis

We performed differential gene expression (DGE) analysis to evaluate transcriptional changes across conditions for each cell line. Variability in the number of differentially expressed genes (DEGs) was observed across batches, cell lines, and conditions. Histograms (Figure 4) illustrate significant differences in DEG counts, with HepG2 cells consistently showing higher numbers of DEGs than HEK293 and HeLa cells.

**Figure 4.**
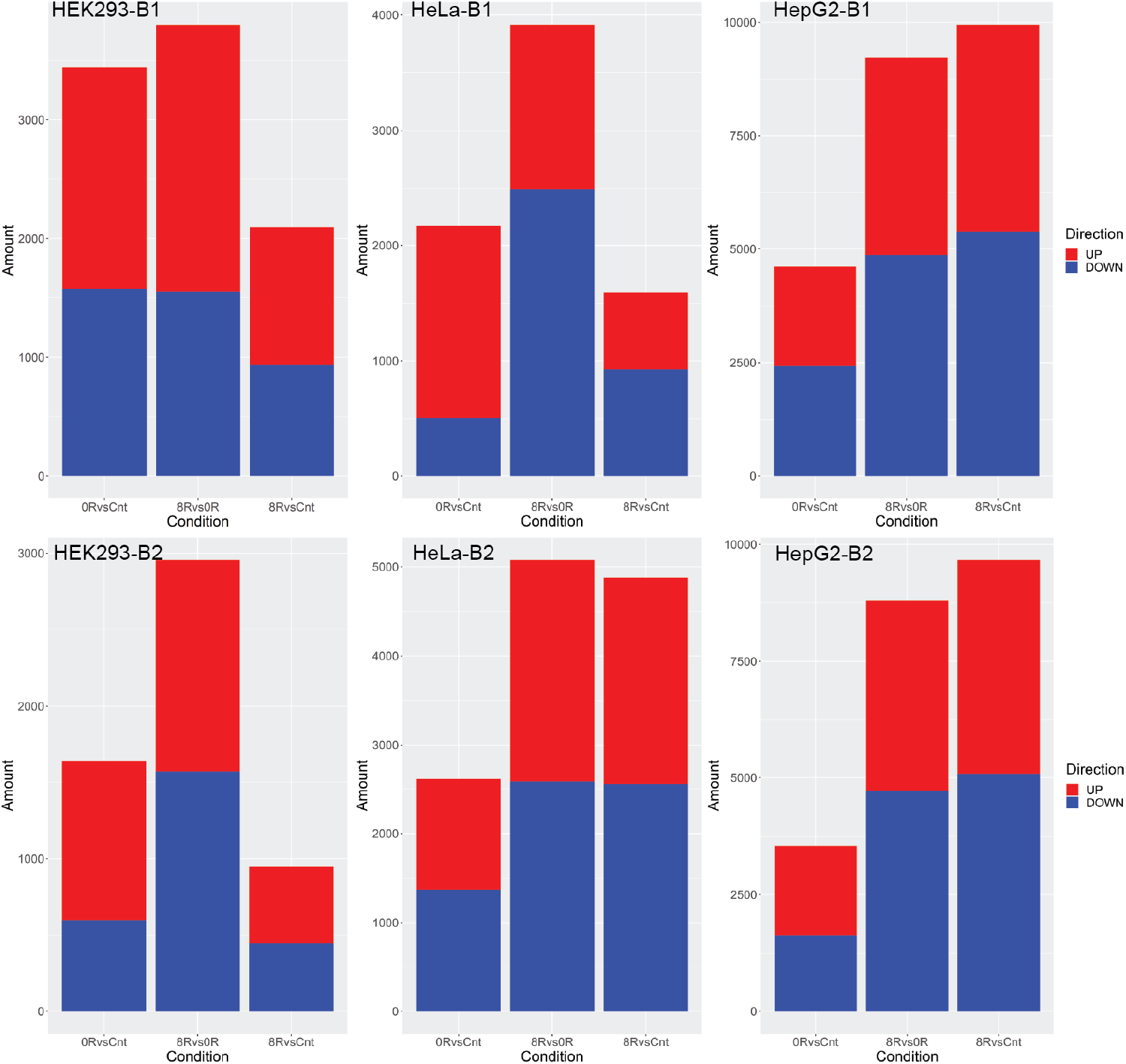
The Number of Differentially Expressed Genes varies by Cell Line and Batch. Histogram depicting the number of differentially expressed genes (|log2 fold change| > 0.5; adjusted p-value < 0.05) in HEK293 (Figures 4A, 4B), HeLa (Figures 4C, 4D), and HepG2 (Figures 4E, 4F) cell lines during heat shock for Batch 1 (left) and Batch 2 (right). The height of each bar represents the count of upregulated and downregulated genes, emphasizing the scale of transcriptional changes in response to heat shock. Note that the Y-axis scale varies across panels to improve visualization of the differences in gene expression dynamics between cell lines and batches.

For Batch 1 (Figure 4 top panels), the DEG counts for HEK293 were 3,444 (0 hours vs. Control), 2,096 (8 hours vs. Control), and 3,802 (8 hours vs. 0 hours). Similarly, HeLa showed 2,168, 1,590, and 3,916 DEGs for these comparisons, respectively, while HepG2 exhibited 4,609, 9,948, and 9,219 DEGs. In Batch 2 (Figure 4 bottom panels), HEK293 had 1,639, 947, and 2,955 DEGs; HeLa had 2,613, 4,877, and 5,076 DEGs; and HepG2 exhibited 3,547, 9,665, and 8,791 DEGs for the exact comparisons.

Volcano plots (Figure 5 and Supplementary Figure 2) visualized DEGs’ fold change and statistical significance across comparisons. This approach highlights the fold change and statistical significance of gene expression for each comparison: in HEK293, 1,867 genes were overexpressed, and 1,577 genes were underexpressed in the 0hrvsCnt condition for Batch 1. In HeLa, 1,661 genes were overexpressed, and 507 were underexpressed in the 0hrvsCnt condition for Batch 1. In HepG2, 2,188 genes were overexpressed, and 2,421 genes were underexpressed in the 0hrvsCnt condition for Batch 1. Key heat shock response genes, including *HSPA6, HSPA1A, HSPA1B, BAG3*, and *IER5*, were significantly overexpressed across all cell lines and conditions, with HepG2 cells demonstrating the most robust transcriptional response.

**Figure 5.**
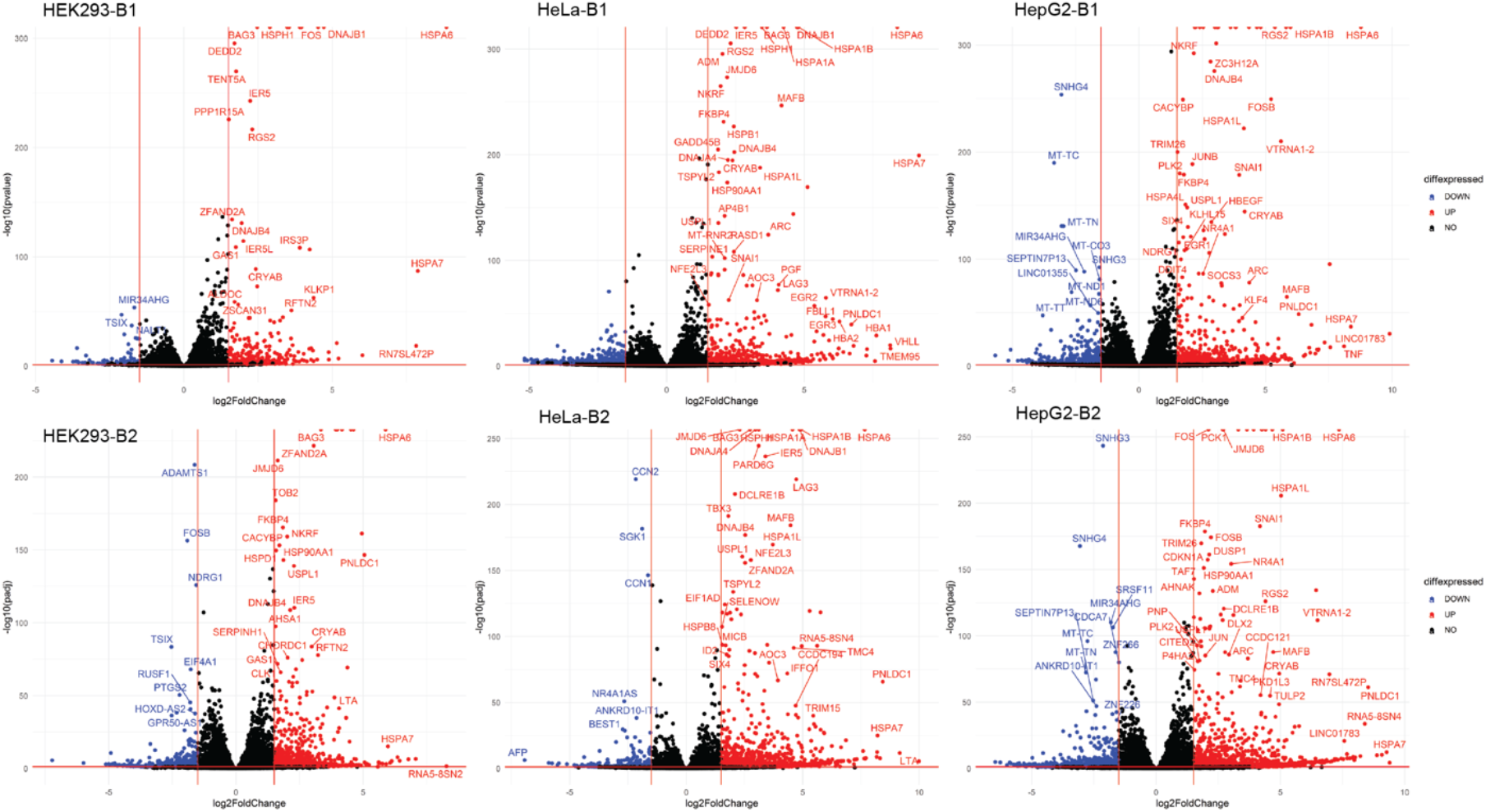
Volcano plots show the distribution of differentially expressed genes for condition comparisons within each cell line. Each point represents a gene, with the x-axis indicating log2 fold change (magnitude of expression change) and the y-axis representing statistical significance (negative log10 adjusted p-value). Genes upregulated (log2FC > 1) are shown in red, while downregulated genes (log2FC < -1) are in blue. Examples include HEK293 (Figure 5A), HeLa (Figure 5B), and HepG2 (Figure 5C) comparisons at 0 hours post-heat shock versus control, visualizing the most significant transcriptional changes.

The next step involved identifying conserved genes shared between the cell lines to further elucidate the similarities in gene expression during mild heat shock across HeLa, HEK, and Hep cells. Conserved DEGs across all three cell lines and batches included 405 genes overexpressed at 0 hours post-heat shock vs. Control, 196 genes overexpressed at 8 hours post-heat shock vs. 0 hours, and 159 genes overexpressed at 8 hours post-heat shock vs. Control (Figures 6A, 6C, and 6E).

**Figure 6.**
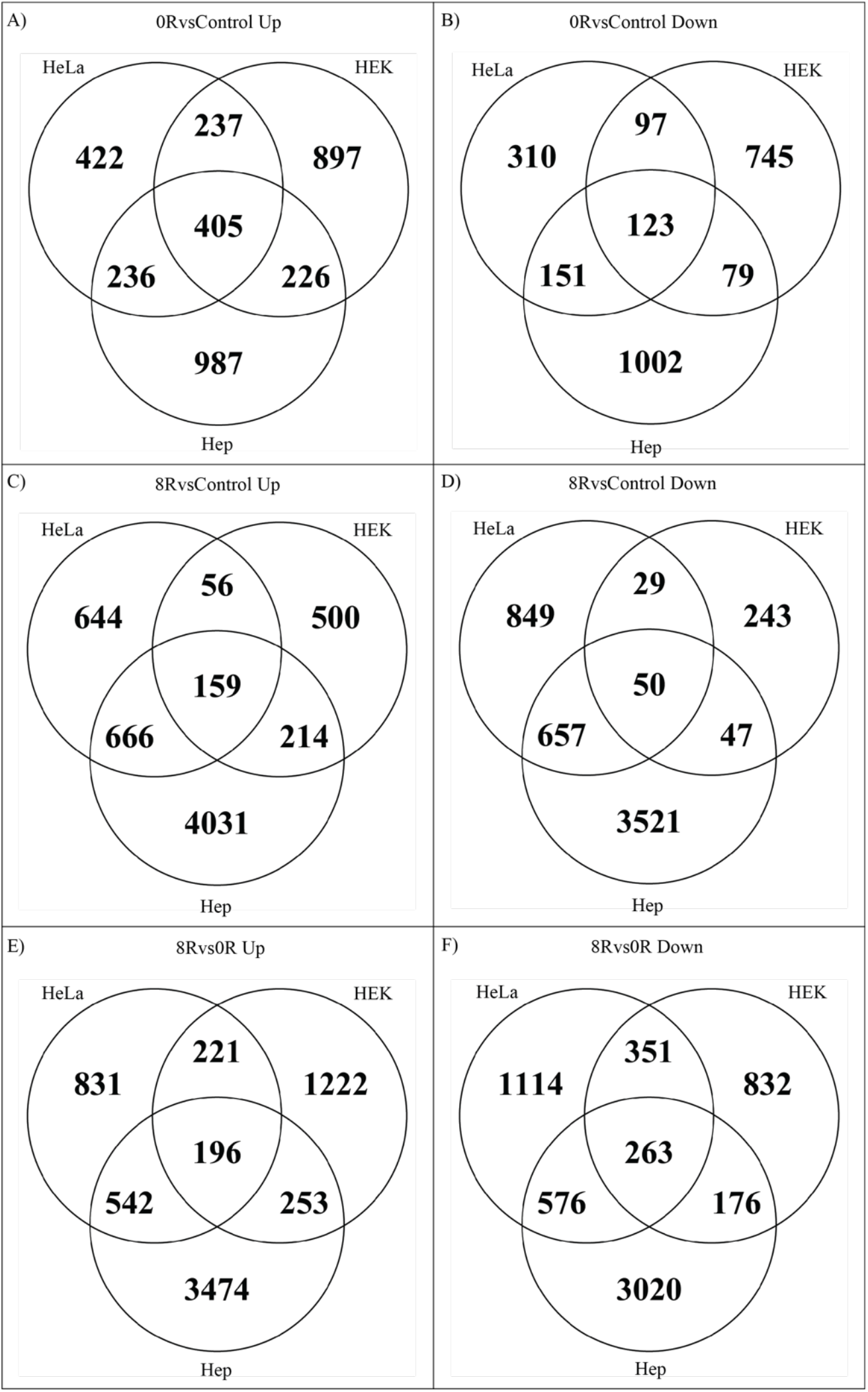
Venn diagrams summarizing conserved differentially expressed genes between cell lines under specific conditions. The size of overlapping circles corresponds to the number of shared genes across comparisons. Overexpressed genes (log2FC > 0.5; p-adj. < 0.05) and underexpressed genes (log2FC < -0.5; p-adj. < 0.05) are visualized for 0 hours post-heat shock vs. control (Figures 6A, 6B), 8 hours vs. 0 hours (Figures 6C, 6D), and 8 hours vs. control (Figures 6E, 6F).

Underexpressed genes shared across all cell lines and batches included 123 genes at 0 hours post-heat shock vs. Control, 263 genes at 8 hours post-heat shock vs. 0 hours, and 50 genes at 8 hours post-heat shock vs. Control (Figures 6B, 6D, and 6F).

### Functional Enrichment Analysis

To elucidate biological processes associated with DEGs, we performed functional enrichment analyses using GSEA and STRING, conducting these analyses separately for each batch due to batch effects. Dot plots (Figure 7 and Supplementary Figure 3) highlight the top ten gene sets with the highest and lowest normalized enrichment scores (NES) across comparisons. In HEK293 (0 hours vs. Control), enriched gene sets related to protein folding and response to unfolded proteins were consistently observed in both batches. However, HeLa and HepG2 showed less overlap in enriched gene sets between batches, suggesting more significant batch-dependent variability.

**Figure 7.**
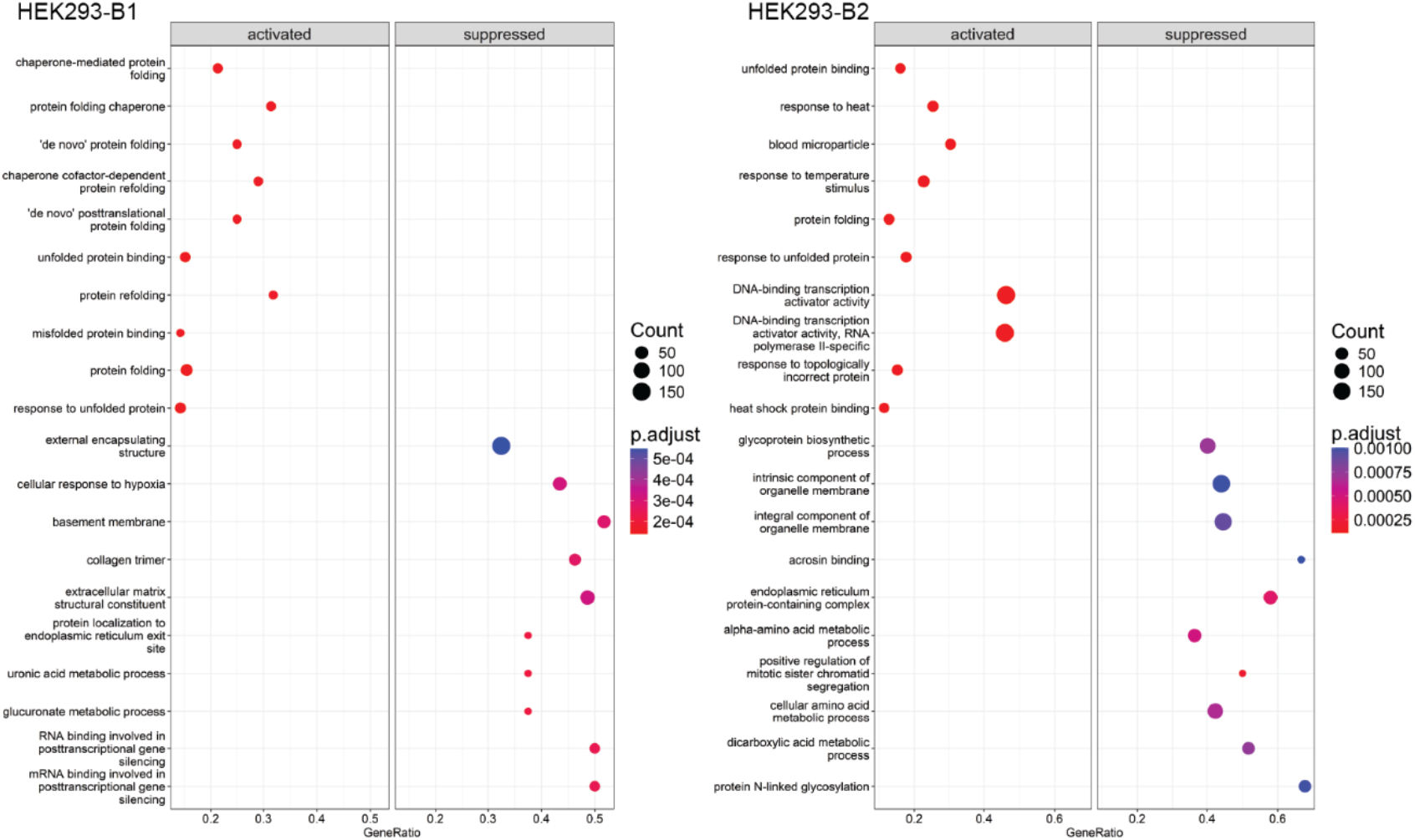
Dot plot visualization of Gene Set Enrichment Analysis (GSEA) results. The top 15 positively and negatively enriched gene sets, ranked by normalized enrichment scores (NES), are shown for HEK293 at 0 hours post-heat shock vs. control (Batch 1 and Batch 2). Each dot represents a gene set, with size proportional to the number of enriched genes and color reflecting the statistical significance (adjusted p-value).

Distribution plots (Figure 8 and Supplemental Figure 4) of enriched genes within the top 15 NES gene sets revealed similar enrichment profiles for genes involved in protein folding and chaperone activity in HEK293. In contrast, unique enrichment distributions were observed in HeLa and HepG2, reflecting differences in stress response pathways.

**Figure 8.**
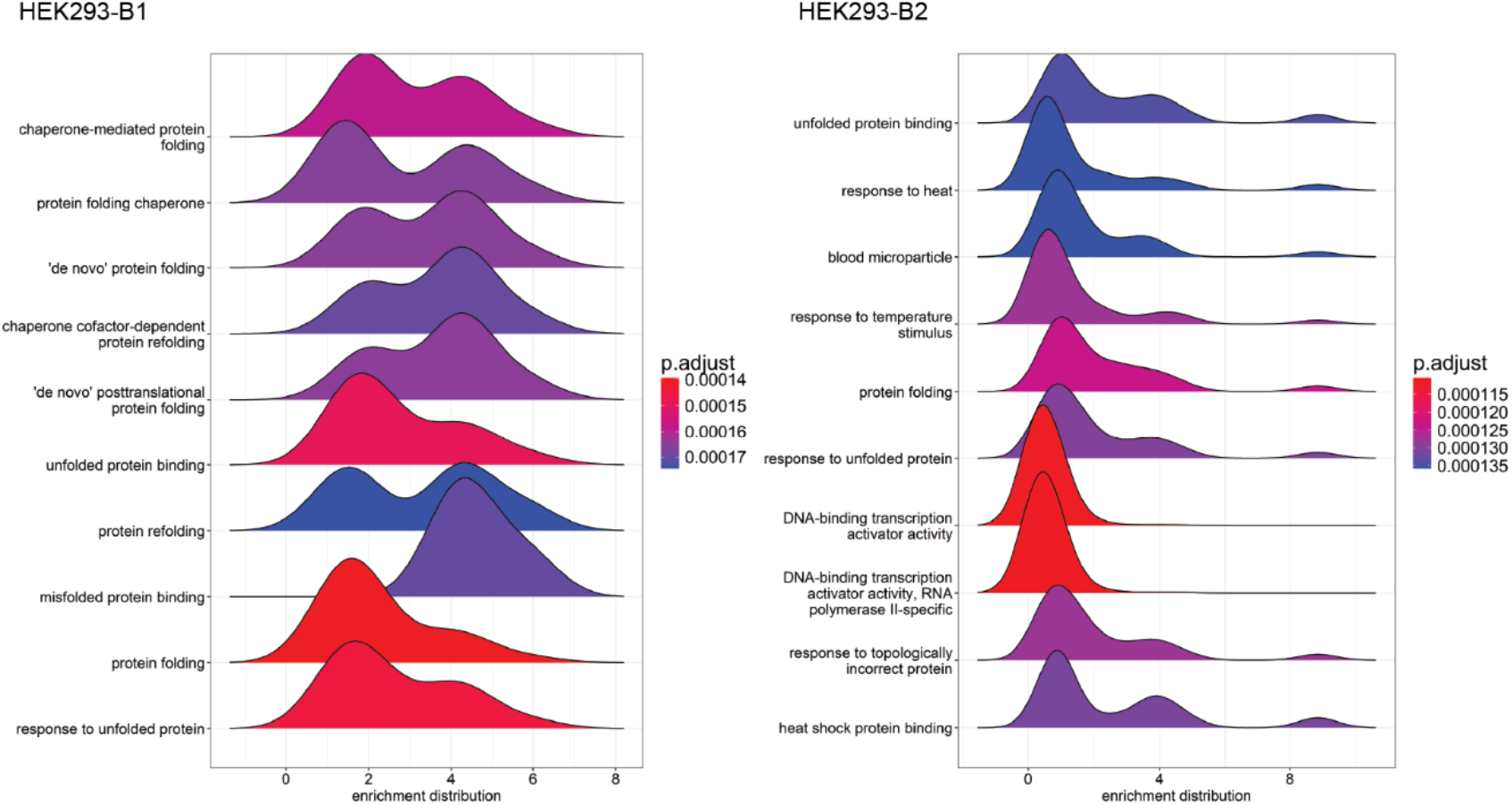
Distribution of Log2 Fold Changes in Enriched Gene Sets. Distribution of log2 fold changes for the top 15 positively enriched GSEA hits in HEK293, HeLa, and HepG2 cell lines at 0 hours post-heat shock vs. control. The peak height represents the number of genes within each range of log2 fold change, highlighting the degree of enrichment within gene sets. Separate plots depict Batch 1 and 2 results for each cell line (e.g., Figures 8A–8F).

Enrichment maps (Figure 9 and Supplementary Figures 5C–5F) visualized interactions among enriched gene sets. In HEK293, gene sets related to protein folding and heat shock response formed cohesive clusters. In HeLa and HepG2, fewer interconnections were observed, with unexpected enrichments in immune-related pathways, such as B cell-mediated immunity and immunoglobulin receptor binding, in HepG2.

**Figure 9.**
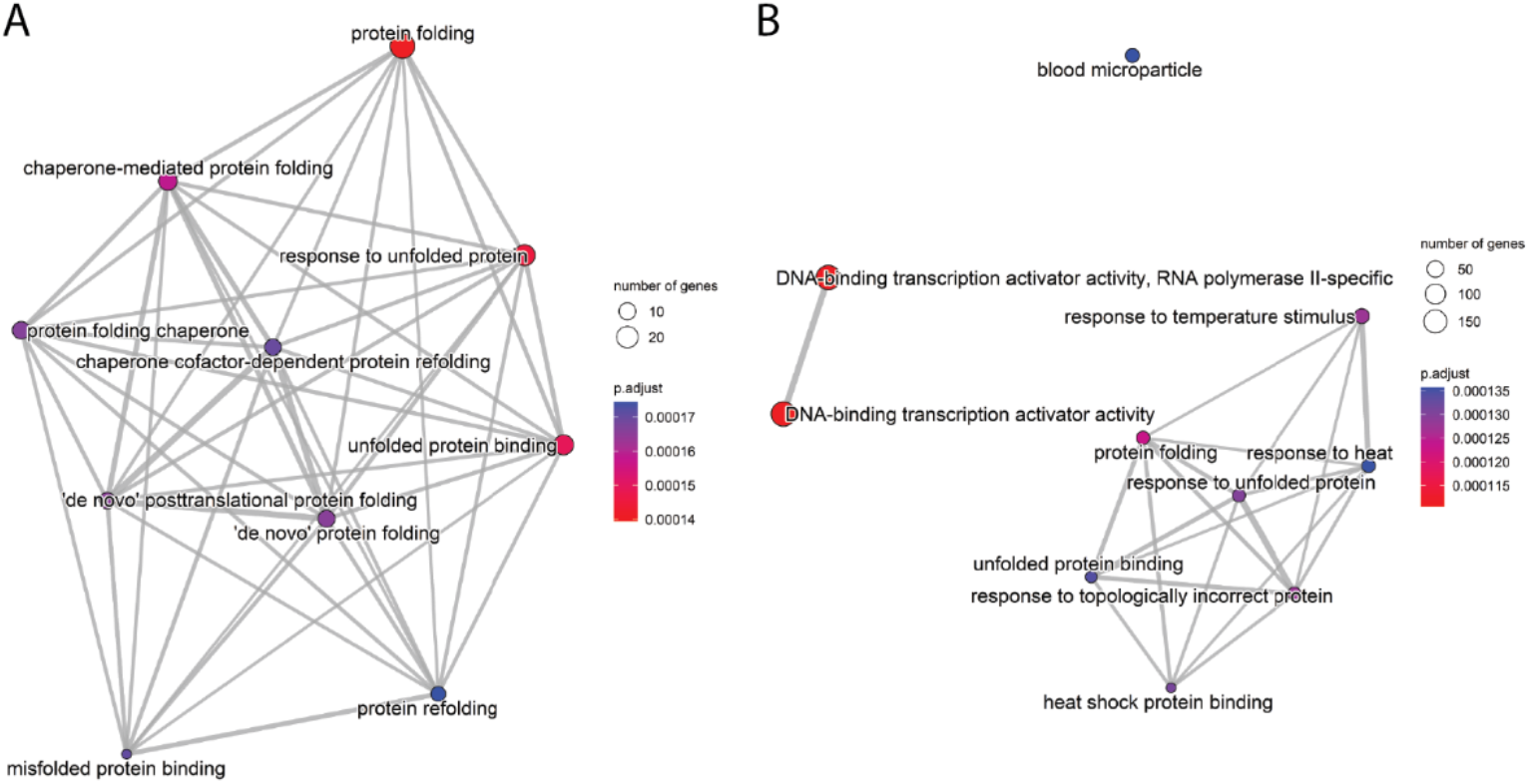
Enrichment maps of the top 10 positively enriched gene sets (GSEA NES) for HEK293 cells at 0 hours post-heat shock vs. control. Each node represents a gene set, with edges indicating shared genes between sets. The network visualizes functional overlap and relationships among the most significant pathways for each batch (Figures 9A, 9B).

STRING analysis identified conserved gene sets enriched across all cell lines and conditions. A total of 18 STRING profiles were generated (three cell lines, three conditions, two batches). Compared to the control sample, four gene sets were enriched and shared between HEK293, HepG2, and HeLa cell lines immediately following mild heat shock. These gene sets are Receptor Ligand Activity (GO:0048018), Signaling Receptor Activator Activity (GO:0030545), Protein Folding Chaperone (GO:0044183), and Class A/1 Rhodopsin-like Receptors (HSA-373076). Notably, the *Receptor Ligand Activity* pathway (GO:0048018) was consistently enriched across all conditions, cell lines, and batches (Table 1).

**Table 1.** Enriched gene sets were conserved in both batches and HEK293, HepG2, and HeLa cell lines.

| 0RvsControl |  | 8RvsCnt |  | 8Rvs0R |  |
| --- | --- | --- | --- | --- | --- |
| ID | Name | ID | Name | ID | Name |
| GO:0048018 | Receptor Ligand Activity | GO:0048018 | Receptor Ligand Activity | GO:0048018 | Receptor Ligand Activity |
| GO:0030545 | Signaling receptor activator activity | hsa04080 | Neuroactive ligand-receptor interaction | hsa04080 | Neuroactive ligand-receptor interaction |
| GO:0044183 | Protein folding chaperone | HSA-500792 | GPCR ligand binding | HSA-500792 | GPCR ligand binding |
| HSA-373076 | Class A/1 (Rhodopsin-like receptors) |  |  | HSA-373076 | Class A/1 (Rhodopsin-like receptors) |

### Analysis of Receptor Ligand Activity (GO:0048018) Genes

Of the 75 genes associated with the *Receptor Ligand Activity* pathway, 13 were consistently expressed across both batches and all three cell lines under each condition. These genes, including *TNF, GNRH2, PSPN, MIA, SEMA4D*, and *HBEGF*, are involved in cell survival, repair, and growth processes (Figure 10). Interaction networks (Figure 11) further illustrate how these genes interact to coordinate stress responses. As anticipated, the gene expression patterns of the conserved genes are similar between the cell lines and batches throughout heat shock.

**Figure 10.**
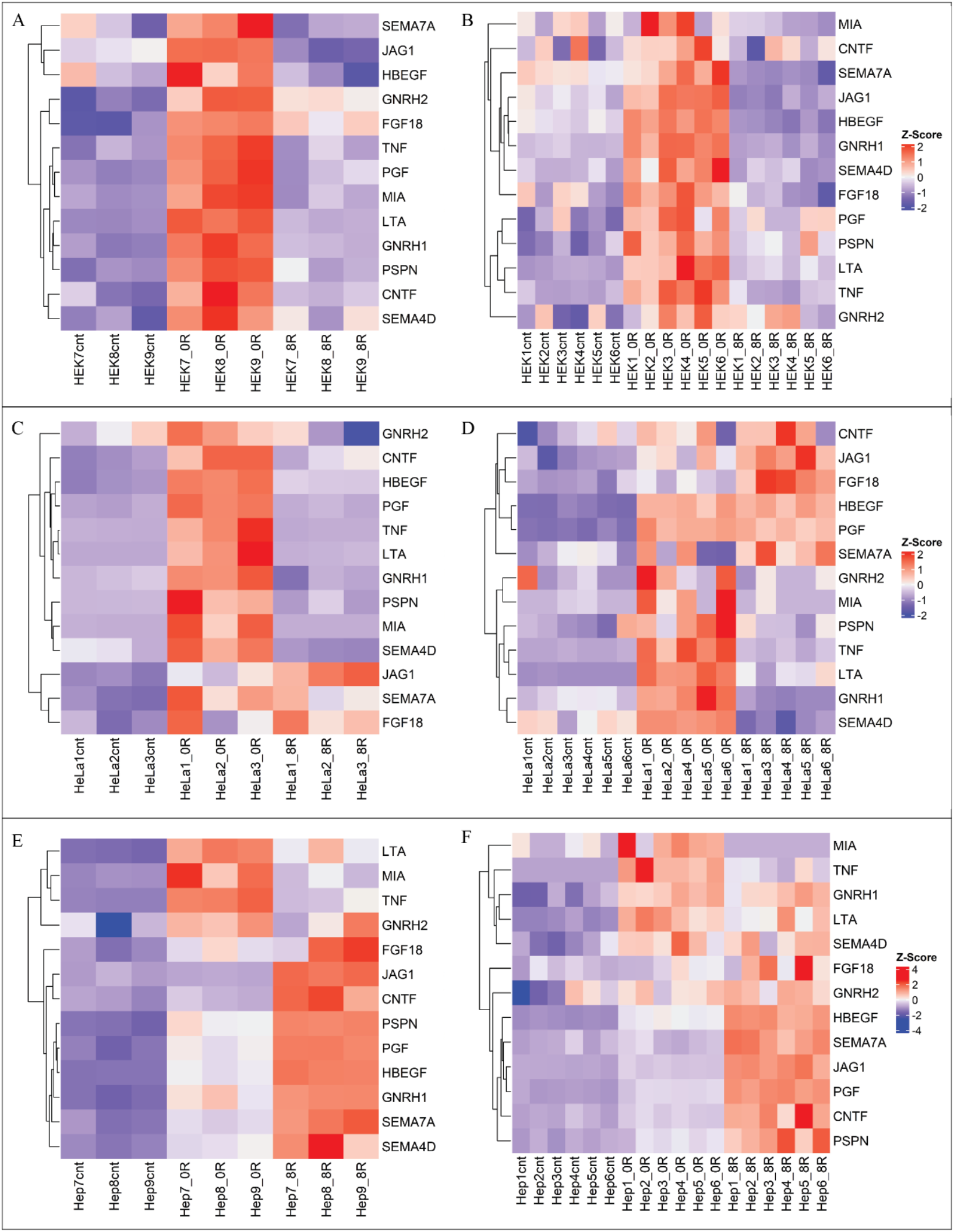
Distribution of Log2 Fold Changes in Enriched Gene Sets. Heatmaps of Z-score-scaled log2 fold changes for 13 conserved genes within the Signal Receptor Ligand Activity (GO:0048018) pathway. Rows represent genes, while columns correspond to cell lines and conditions. Separate heatmaps are provided for Batch 1 (Figures 10A, 10C, 10E) and Batch 2 (Figures 10B, 10D, 10F), illustrating consistent patterns of gene regulation across batches and cell lines.

**Figure 11.**
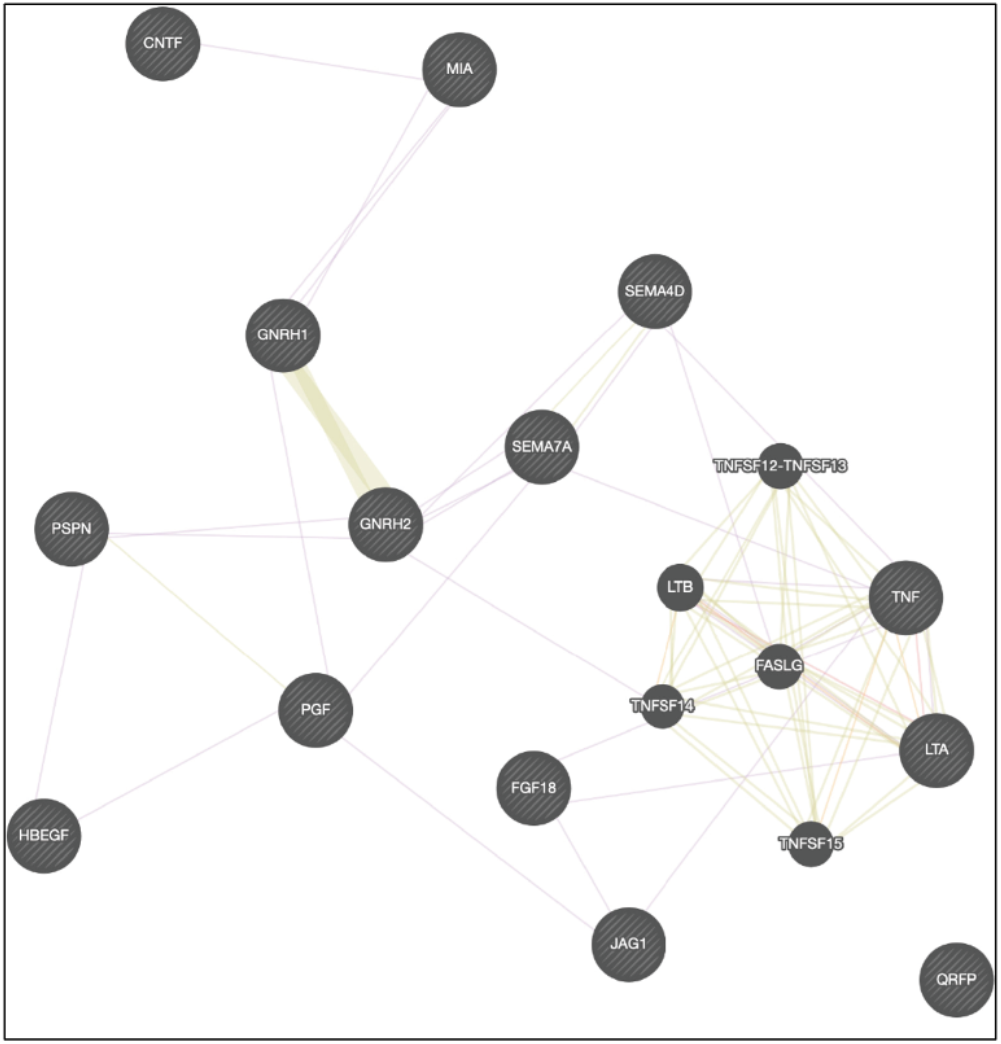
Network Map of GO:0048018 Pathway Genes. Cytoscape-generated network map of the 13 conserved differentially expressed genes within the Signal Receptor Ligand Activity (GO:0048018) pathway. Nodes represent genes, and edges indicate predicted functional interactions, offering a systems-level view of pathway dynamics.

However, some variations in the magnitude of expression can be observed. Supplementary Table 4 provides an overview of the biological processes associated with these 13 genes, many promoting cell survival, repair, and growth.

### Gene Expression Assessment via qPCR

We conducted qPCR on selected genes to validate the differential gene expression (DGE) analysis findings. This approach confirmed our transcriptomic results and provided additional insight into the induction of the heat shock response. The *Heat Acclimation* gene set (GO:0010286) was chosen for its relevance, containing six genes known to exhibit significant expression changes during heat shock.

Figure 12 summarizes the expression changes in HEK293 (Figures 12A, 12B), HeLa (Figures 12C, 12D), and HepG2 (Figures 12E, 12F) cell lines across batches and conditions. These results strongly support the activation of the heat shock response. *HSPA1A, HSPA1B*, and *HSPA6* were consistently upregulated across all cell lines, while *RBBP7* was downregulated. However, expression patterns of *HSBP1* and *HSBP1L1* varied. In HEK293 cells, *HSBP1* decreased at 0- and 8-hours post-heat shock, whereas it increased in HeLa and HepG2 cells under the same conditions. Conversely, *HSBP1L1* increased in HEK and HeLa cells but exhibited batch-specific behavior in HepG2, decreasing in Batch 1 at 8 hours and remaining unchanged in Batch 2. Despite this variability, the consistent expression trends of *HSPA1A, HSPA1B, HSPA6*, and *RBBP7* across all conditions underscore the canonical activation of the heat shock response.

**Figure 12.**
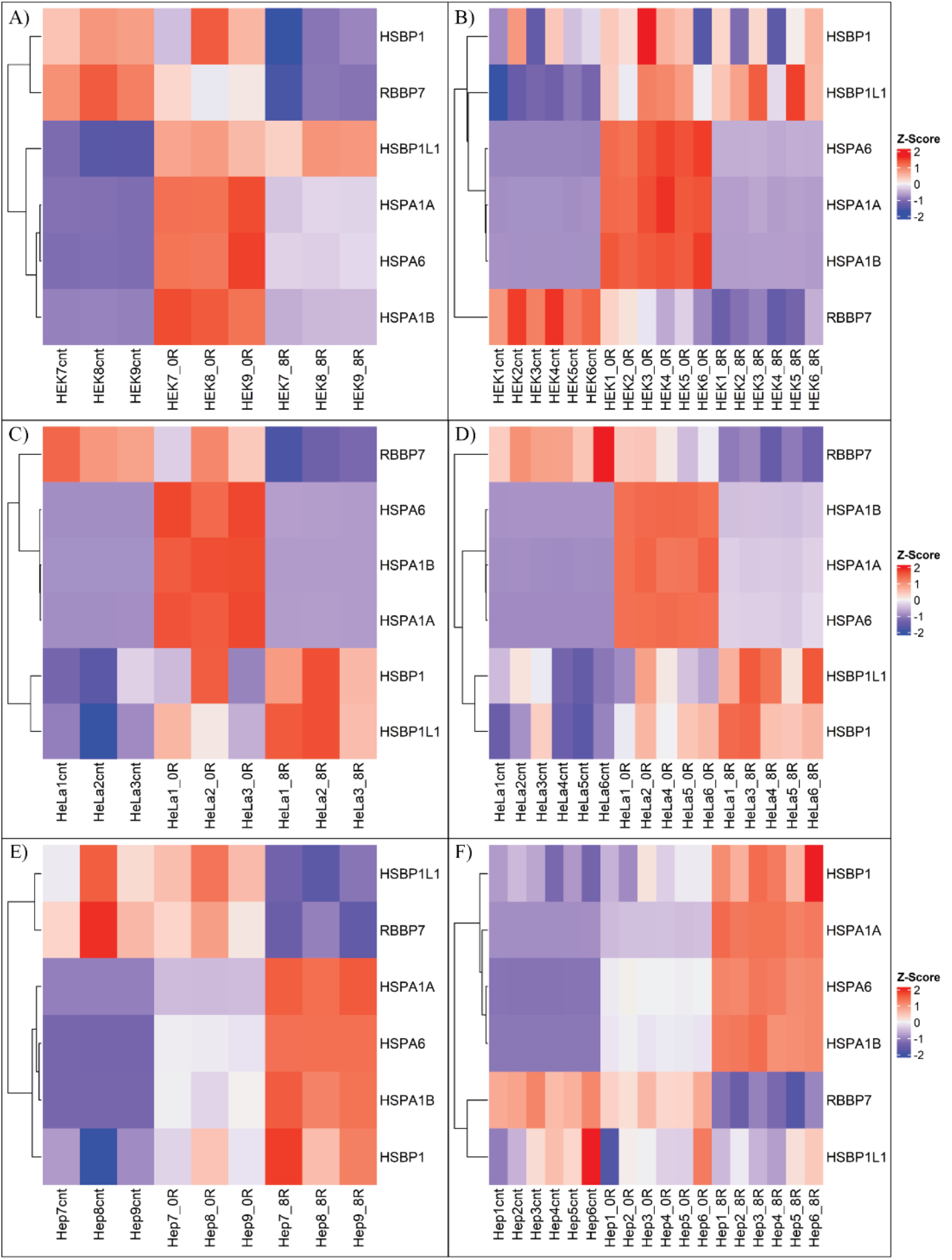
Heatmaps of Heat Acclimation Genes Across Conditions. Heat maps show expression changes for Heat Acclimation (GO:0010286) genes at control, 0 hours post-heat shock, and 8 hours post-heat shock conditions. Separate heatmaps are provided for Batch 1 (Figures 12A, 12C, 12E) and Batch 2 (Figures 12B, 12D, 12F) across HEK293, HeLa, and HepG2 cell lines.

qPCR analysis was further performed on mRNA from HeLa cells at 0- and 8-hours post-heat shock, along with control samples maintained at 37°C. The primers targeted four “Heat Acclimation” genes (*HSPA1A, HSPA6, BAG3, DNAJB1*) and three “Signal Receptor Ligand Activity” genes (*LTA, MIA, TNF*), with *ACTB* and *GAPDH* as reference controls.

As shown in Figure 13, *HSPA1A* and *HSPA6* exhibited the highest fold changes at 0- and 8 hours post-heat shock (Figure 13A), aligning with their central roles in the heat shock response. *BAG3* and *DNAJB1* also displayed significant changes in expression (Figure 13B). For the “Signal Receptor Ligand Activity” genes, *LTA, MIA*, and *TNF* were upregulated immediately after heat shock but returned to near-control levels by 8 hours post-recovery (Figure 13C). The observed patterns closely mirrored those revealed by RNA-seq analyses, except HSPA1A and HSPA6, which continued to increase at 8 hours post-recovery in the qPCR data. Although the log2 fold change values varied slightly between the two methods, the overall trends were consistent (Supplemental Figure 6).

**Figure 13.**
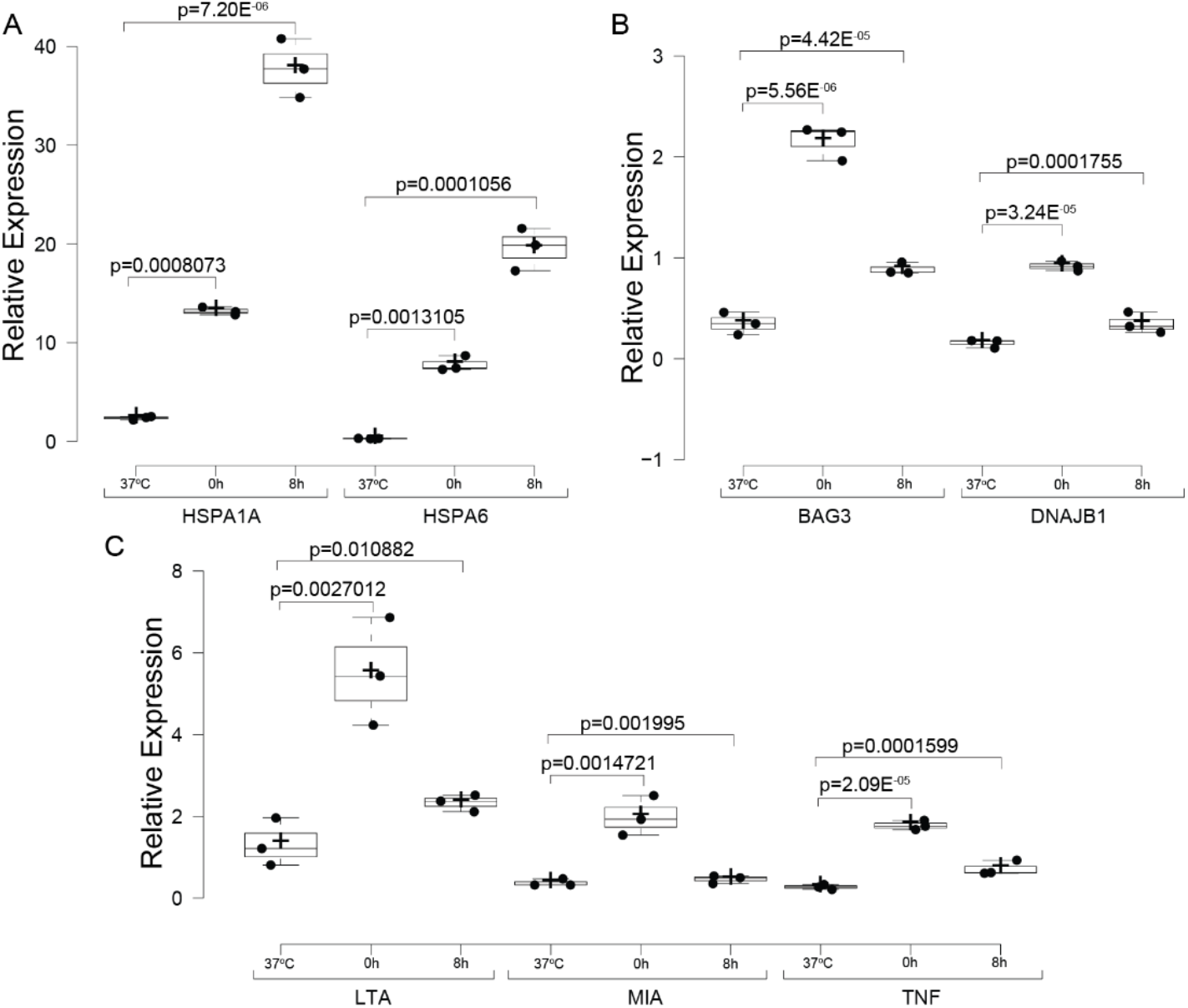
qPCR Validation of Key Gene Sets. mRNA fold change measured by qPCR for Heat Acclimation (GO:0010286) and Signal Receptor Ligand Activity (GO:0048018) genes in HeLa cells at control, 0 hours post-heat shock, and 8 hours post-heat shock. Results are normalized against housekeeping genes (ACTB and GAPDH). Heat shock genes (HSPA1A, HSPA6) exhibit the highest induction, while signal receptor ligand genes (LTA, MIA, TNF) show transient upregulation (Figures 13A–13C). Note that the Y-axis scale varies across panels. Statistical significance was determined using one-way ANOVA (Analysis of Variance) followed by post-hoc Tukey HSD (Honestly Significant Difference) and Bonferroni tests.

These results validate the involvement of key “Heat Acclimation” and “Signal Receptor Ligand Activity” genes in the heat shock response. The consistency between qPCR findings and DGE and functional enrichment analyses reinforces the robustness of our conclusions and highlights specific pathways central to stress adaptation mechanisms.

## Discussion

This study characterizes the heat shock response (HSR) in three human cell lines—HeLa, HEK293, and HepG2—by subjecting them to mild heat shock and examining recovery periods of 0 and 8 hours. Using principal component analysis (PCA), differential gene expression (DGE), and functional enrichment analyses, we identified conserved and cell-specific genes and pathways activated during and after heat shock. PCA revealed that the primary variance in gene expression was driven by cell type, batch effects, and heat shock recovery time points (Figures 2–3). To address the loss of statistical power caused by splitting into two experimental batches, we analyzed the batches individually, effectively gaining a second independent repetition and ensuring that observed trends were consistent across both sets of experiments. Collectively, these findings confirm the activation of canonical HSR pathways while also highlighting novel pathways, including “Receptor Ligand Activity” (GO:0048018), which were consistently enriched across all cell lines and conditions (Table 1, Figures 10–11).

While the heat shock response was conserved across all cell lines, the transcriptional responses revealed cell-type-specific adaptations, particularly in cancerous versus non-cancerous cells. The observed activation of well-characterized heat shock protein (HSP) genes, including *HSPA1A, HSPA1B*, and *HSPA6*, underscores their central role in managing proteotoxic stress (Figures 5, 9, and 12). In HEK293 cells, we observed a strong upregulation of genes involved in protein folding and chaperone-mediated activities, particularly during the 0-hour recovery period. This reflects the robustness of the HSR in these cells to mitigate misfolded protein accumulation and restore homeostasis [1-13]. However, it is important to note that these experiments were conducted in established cell lines, which, while offering precise control over experimental variables, lack the complexity and tissue-level interactions present in animal models [37].

In contrast, the transcriptional responses in cancerous HeLa and HepG2 cells were more dynamic, particularly at the 8-hour recovery stage. For example, over 9,500 genes were differentially expressed in HepG2 cells after 8 hours of recovery, reflecting a heightened transcriptional activity likely driven by their reliance on enhanced proteostasis mechanisms to sustain rapid growth and manage misfolded proteins under adverse conditions [38-41]. Despite these differences, core stress response mechanisms were conserved across all cell lines. This highlights the universal importance of fundamental pathways like protein folding and chaperoning in maintaining cellular homeostasis after stress. The shared upregulation of HSPs and related pathways underscores their centrality in preventing apoptosis and sustaining proteostasis under heat-induced stress [2,38].

Indeed, cancer cells such as HeLa and HepG2 demonstrated a more extensive and dynamic heat shock response than non-cancerous HEK293 cells. This aligns with the chronic stress conditions encountered by cancer cells—such as hypoxia, oxidative damage, and immune evasion—that make their survival heavily dependent on stress response pathways like the unfolded protein response (UPR) and heat shock proteins (HSPs) [38,40]. The marked increase in HSP expression, particularly HSPA1A, HSPA1B, and HSPA6, highlights their dual role in managing protein misfolding and protecting against apoptosis (Figures 5, 9, and 12). These heightened stress responses likely contribute to cancer cell survival under hostile conditions, including during therapy. For example, we observed significant transcriptional shifts in HepG2 cells at 8 hours post-heat shock, with substantial upregulation of HSR-related genes [1,38].

These findings point to potential mechanisms through which stress signaling pathways are leveraged in cells, including cancer cells, to enhance survival under stress. Indeed, a notable finding of this study was the consistent enrichment of the “Receptor Ligand Activity” pathway (GO:0048018) across all conditions and cell lines (Table 1, Figures 10–12). Traditionally associated with extracellular signaling, its activation during heat shock suggests a novel role in mediating cellular responses to proteotoxic stress. Receptor-ligand interactions may serve as conduits for communicating stress signals, triggering adaptive responses such as apoptotic regulation or survival signaling [42]. In multicellular systems, these pathways may synchronize stress responses across cell populations to preserve tissue integrity, balancing survival, and apoptosis to influence outcomes following stress exposure [43,44]. Further exploration of the receptor-ligand pairs involved in the HSR could uncover new stress adaptation and resilience mechanisms.

The consistent activation of the “Receptor Ligand Activity” pathway in cancerous and non-cancerous cells underscores its potential role in stress resilience. In cancer cells, receptor-ligand interactions may help coordinate stress signals within the tumor microenvironment, enhancing collective survival under stress [42]. Identifying receptors uniquely or commonly expressed on the surfaces of HeLa, HepG2, and HEK293 cells could provide valuable insights into how receptor-ligand signaling mediates the HSR. Receptors conserved across all cell lines or specific to cancer cells could represent likely candidates for immediate regulation by the HSR, either following heat shock or during recovery. Targeting receptor-ligand signaling could offer a novel therapeutic strategy by disrupting stress communication networks, complementing existing therapies such as heat shock protein inhibitors to reduce tumor resilience to proteotoxic stress.

## Conclusion

In conclusion, this study highlights the conserved nature of cellular stress responses across diverse cell types and the unique adaptations cancer cells employ to survive under stress. The consistent activation of “Receptor Ligand Activity” during heat shock offers a novel avenue for exploring stress communication mechanisms and their role in cancer resilience. The upregulation of key HSPs in cancer cells underscores their centrality in stress adaptation, making them attractive therapeutic targets. These findings provide valuable insights into the HSR’s conserved and cancer-specific aspects, laying the groundwork for developing interventions that exploit the vulnerabilities of stress-adapted cells.

## Supporting information

DEG files

## Supplementary Materials

The manuscript includes supplementary figures and tables.

**Appendix 1** Contains the complete detailed methodology used in the transcriptomics analysis.

**Appendix 2** Contains the DEG tables for all comparisons of this study (with a Log2FC >0.5 and p. adj. <0.05).

## Author Contributions

Conceptualization, Nikolas Nikolaidis; Data curation, Andrew Reinschmidt and Nikolas Nikolaidis; Formal analysis, Andrew Reinschmidt, Luis Solano, Yonny Chavez, William Drew Hulsy, and Nikolas Nikolaidis; Funding acquisition, Nikolas Nikolaidis; Methodology, Andrew Reinschmidt, Luis Solano, and Nikolas Nikolaidis; Resources, Nikolas Nikolaidis; Writing – original draft, Andrew Reinschmidt and Nikolas Nikolaidis; Writing – review & editing, Andrew Reinschmidt, Luis Solano, Yonny Chavez, William Drew Hulsy, and Nikolas Nikolaidis.

## Funding

The research reported in this publication was supported by the National Institute of General Medical Sciences of the National Institutes of Health under Award Number SC3GM121226. The content is solely the authors’ responsibility and does not necessarily represent the official views of the National Institutes of Health.

### Data Availability Statement

All data reported are provided in the text, supplemental figures, or Appendix 1 and 2. The raw data are hosted at NCBI (GEO). The scripts used can be found at (https://github.com/ajreinschmidt/MSc-Code/blob/main/iSEE%20Analysis%20Script.R)

## Acknowledgments

We thank Dr. Dimitra Chalkia for her valuable comments and help with the analysis of the manuscript.

