## supplemental figures and Tables for "Transcriptomics Unveil Canonical and Non-Canonical Heat Shock-Induced Pathways in Human Cell Lines"

**Title**

**Supplementary Table 1.** Summary table of quality control for Batch 1.

| Sample name | Raw reads | Clean reads | Raw bases | Clean bases | Error rate(%) | Q20(%) | Q30(%) | GC content(%) |
|---|---|---|---|---|---|---|---|---|
| HeLa1cnt | 35872594 | 35392491 | 10.8G | 10.6G | 0.03 | 97.86 | 94.18 | 50.52 |
| HeLa2cnt | 40588307 | 39763626 | 12.2G | 11.9G | 0.03 | 97.76 | 93.96 | 50.28 |
| HeLa3cnt | 48968978 | 47830612 | 14.7G | 14.3G | 0.03 | 97.85 | 94.19 | 50.58 |
| HeLa1_0R | 31967532 | 31443462 | 9.6G | 9.4G | 0.03 | 97.69 | 93.84 | 50.66 |
| HeLa2_0R | 37072811 | 36192382 | 11.1G | 10.9G | 0.02 | 97.89 | 94.33 | 50.42 |
| HeLa3_0R | 41466226 | 40777276 | 12.4G | 12.2G | 0.03 | 97.8 | 94.11 | 50.71 |
| HeLa1_8R | 30312159 | 29652893 | 9.1G | 8.9G | 0.03 | 97.76 | 93.96 | 51.29 |
| HeLa2_8R | 35227814 | 34635299 | 10.6G | 10.4G | 0.03 | 97.86 | 94.2 | 50.45 |
| HeLa3_8R | 33843797 | 33328748 | 10.2G | 10.0G | 0.03 | 97.86 | 94.21 | 49.7 |
| HEK1cnt | 39926163 | 39126229 | 12.0G | 11.7G | 0.03 | 97.79 | 94.06 | 48.43 |
| HEK2cnt | 34041857 | 33416198 | 10.2G | 10.0G | 0.03 | 97.78 | 94.03 | 48.92 |
| HEK3cnt | 35956039 | 35315742 | 10.8G | 10.6G | 0.03 | 97.73 | 93.89 | 48.31 |
| HEK1_0R | 40629763 | 40099360 | 12.2G | 12.0G | 0.03 | 97.61 | 93.7 | 48.2 |
| HEK2_0R | 38002642 | 37410561 | 11.4G | 11.2G | 0.03 | 97.52 | 93.5 | 48.3 |
| HEK3_0R | 40373617 | 39155548 | 12.1G | 11.7G | 0.03 | 97.72 | 93.96 | 49.14 |
| HEK1_8R | 33442598 | 32604721 | 10.0G | 9.8G | 0.03 | 97.68 | 93.86 | 48.38 |
| HEK2_8R | 35327546 | 34572374 | 10.6G | 10.4G | 0.03 | 97.82 | 94.22 | 48.03 |
| HEK3_8R | 42653168 | 41520036 | 12.8G | 12.5G | 0.03 | 97.67 | 93.83 | 48.5 |
| Hep1cnt | 35856136 | 34755018 | 10.8G | 10.4G | 0.03 | 97.71 | 93.84 | 48.92 |
| Hep2cnt | 32429635 | 31241941 | 9.7G | 9.4G | 0.03 | 97.73 | 93.84 | 45.92 |
| Hep3cnt | 38344302 | 37715012 | 11.5G | 11.3G | 0.03 | 97.7 | 93.87 | 48.28 |
| Hep1_0R | 42554874 | 41628733 | 12.8G | 12.5G | 0.03 | 97.55 | 93.67 | 50.26 |
| Hep2_0R | 38212434 | 37586682 | 11.5G | 11.3G | 0.03 | 97.61 | 93.69 | 49.57 |
| Hep3_0R | 39235855 | 38542215 | 11.8G | 11.6G | 0.03 | 97.56 | 93.66 | 50.12 |
| Hep1_8R | 32004931 | 31142253 | 9.6G | 9.3G | 0.03 | 97.52 | 93.54 | 47.79 |
| Hep2_8R | 43290892 | 42319750 | 13.0G | 12.7G | 0.03 | 97.36 | 93.14 | 48.29 |
| Hep3_8R | 38439567 | 37525468 | 11.5G | 11.3G | 0.03 | 97.43 | 93.33 | 48.47 |

**Supplementary Table 2.** Summary table of quality control for Batch 2.

| Sample name | Raw reads | Clean reads | Raw bases | Clean bases | Error rate (%) | Q20(%) | Q30(%) | GC content (%) |
|---|---|---|---|---|---|---|---|---|
| HeLa1cnt | 62016498 | 9.3G | 60845478 | 9.13G | 0.02 | 98.21 | 94.87 | 49.5 |
| HeLa2cnt | 70595362 | 10.59G | 69120548 | 10.37G | 0.02 | 98.23 | 95.01 | 49.52 |
| HeLa3cnt | 60631278 | 9.09G | 59517000 | 8.93G | 0.02 | 98.23 | 95.13 | 50.53 |
| HeLa4cnt | 81419890 | 12.21G | 80080716 | 12.01G | 0.02 | 98.19 | 94.82 | 49.28 |
| HeLa5cnt | 73492284 | 11.02G | 72283092 | 10.84G | 0.02 | 98 | 94.8 | 49.95 |
| HeLa6cnt | 62844008 | 9.43G | 61885638 | 9.28G | 0.02 | 98.24 | 94.99 | 49.85 |
| HeLa1_0R | 61676822 | 9.25G | 60913992 | 9.14G | 0.02 | 98.28 | 95.18 | 50.3 |
| HeLa2_0R | 61288296 | 9.19G | 60459190 | 9.07G | 0.02 | 98.31 | 95.23 | 50.05 |
| HeLa4_0R | 63662714 | 9.55G | 62807390 | 9.42G | 0.02 | 98.31 | 95.24 | 49.68 |
| HeLa5_0R | 61124770 | 9.17G | 60088668 | 9.01G | 0.02 | 98.28 | 95.13 | 50.07 |
| HeLa6_0R | 61395698 | 9.21G | 60252900 | 9.04G | 0.02 | 98.26 | 95.19 | 50.69 |
| HeLa1_8R | 61353524 | 9.2G | 60294048 | 9.04G | 0.02 | 98.19 | 94.94 | 50.38 |
| HeLa3_8R | 60850104 | 9.13G | 59765084 | 8.96G | 0.02 | 98.25 | 95.16 | 50.11 |
| HeLa4_8R | 62198866 | 9.33G | 60957704 | 9.14G | 0.02 | 98.02 | 94.48 | 49.66 |
| HeLa5_8R | 63932420 | 9.59G | 62589710 | 9.39G | 0.02 | 98.08 | 94.7 | 50.37 |
| HeLa6_8R | 61521638 | 9.23G | 60228370 | 9.03G | 0.02 | 98.26 | 95.21 | 50.27 |
| HEK1cnt | 66272392 | 9.94G | 65330354 | 9.8G | 0.02 | 98.3 | 95.17 | 49.92 |
| HEK2cnt | 60035250 | 9.01G | 58712308 | 8.81G | 0.02 | 98.23 | 95.07 | 50.36 |
| HEK3cnt | 64145522 | 9.62G | 63225354 | 9.48G | 0.02 | 98.29 | 95.1 | 49.15 |
| HEK4cnt | 65659452 | 9.85G | 64716964 | 9.71G | 0.02 | 98.35 | 95.26 | 48.98 |
| HEK5cnt | 60758956 | 9.11G | 59607368 | 8.94G | 0.02 | 98.26 | 95 | 48.95 |
| HEK6cnt | 61580132 | 9.24G | 60342074 | 9.05G | 0.02 | 98.4 | 95.42 | 49.15 |
| HEK1_0R | 69441930 | 10.42G | 68057050 | 10.21G | 0.02 | 98.07 | 94.72 | 50.09 |
| HEK2_0R | 67217478 | 10.08G | 65897520 | 9.88G | 0.02 | 98.03 | 94.63 | 49.76 |
| HEK3_0R | 61440470 | 9.22G | 60215412 | 9.03G | 0.02 | 98.25 | 95.16 | 48.6 |
| HEK4_0R | 64208408 | 9.63G | 62884812 | 9.43G | 0.02 | 98.26 | 95.24 | 48.42 |
| HEK5_0R | 60640764 | 9.1G | 59344610 | 8.9G | 0.02 | 98.23 | 95.16 | 48.85 |
| HEK6_0R | 60250308 | 9.04G | 59084584 | 8.86G | 0.02 | 98.24 | 95.19 | 49.16 |
| HEK1_8R | 63929742 | 9.59G | 63030320 | 9.45G | 0.02 | 98.42 | 95.47 | 49.5 |
| HEK2_8R | 63883690 | 9.58G | 62590046 | 9.39G | 0.02 | 98.44 | 95.53 | 49.4 |
| HEK3_8R | 78514742 | 11.78G | 76691164 | 11.5G | 0.02 | 98.41 | 95.46 | 48.93 |
| HEK4_8R | 58987108 | 8.85G | 57708896 | 8.66G | 0.02 | 98.29 | 95.14 | 49.43 |
| HEK5_8R | 63175986 | 9.48G | 62004916 | 9.3G | 0.02 | 98.24 | 95.02 | 49.55 |
| HEK6_8R | 63239640 | 9.49G | 62031320 | 9.3G | 0.02 | 98.39 | 95.39 | 49.39 |
| Hep1cnt | 67481398 | 10.12G | 65973190 | 9.9G | 0.02 | 98.4 | 95.42 | 48.33 |
| Hep2cnt | 79463982 | 11.92G | 77844494 | 11.68G | 0.02 | 98.32 | 95.24 | 47.94 |
| Hep3cnt | 62406870 | 9.36G | 61270232 | 9.19G | 0.02 | 98.25 | 95.03 | 48.08 |
| Hep4cnt | 63241240 | 9.49G | 61807164 | 9.27G | 0.02 | 98.24 | 95.04 | 49.3 |

| | | | | | | | | |
|---|---|---|---|---|---|---|---|---|
| Hep5cnt | 65096858 | 9.76G | 63728104 | 9.56G | 0.02 | 98.32 | 95.21 | 48.45 |
| Hep6cnt | 68099944 | 10.21G | 66821570 | 10.02G | 0.02 | 98.19 | 94.86 | 47.62 |
| Hep1_0R | 62107654 | 9.32G | 60784062 | 9.12G | 0.02 | 98.32 | 95.22 | 49.26 |
| Hep2_0R | 60533366 | 9.08G | 59547872 | 8.93G | 0.02 | 98.34 | 95.32 | 49.4 |
| Hep3_0R | 64125286 | 9.62G | 62786732 | 9.42G | 0.02 | 98.32 | 95.22 | 49.1 |
| Hep4_0R | 63549094 | 9.53G | 62115522 | 9.32G | 0.02 | 98.2 | 95.06 | 50 |
| Hep5_0R | 60337954 | 9.05G | 59417138 | 8.91G | 0.02 | 98.17 | 94.86 | 49.09 |
| Hep6_0R | 66796434 | 10.02G | 65204242 | 9.78G | 0.02 | 98.16 | 94.82 | 48.53 |
| Hep1_8R | 65094562 | 9.76G | 63457930 | 9.52G | 0.02 | 98.19 | 94.9 | 48.59 |
| Hep2_8R | 63557536 | 9.53G | 62146416 | 9.32G | 0.02 | 98.15 | 94.85 | 49.11 |
| Hep3_8R | 62153818 | 9.32G | 61130290 | 9.17G | 0.02 | 98.19 | 94.92 | 48.68 |
| Hep4_8R | 64571708 | 9.69G | 62780996 | 9.42G | 0.03 | 97.6 | 93.52 | 48.98 |
| Hep5_8R | 67192788 | 10.08G | 65483534 | 9.82G | 0.03 | 97.69 | 93.66 | 48.87 |
| Hep6_8R | 59336896 | 8.9G | 58500944 | 8.78G | 0.02 | 98.23 | 95.12 | 48.44 |

**Supplementary Table 3.** Primers used in qPCR experiments.

| Classification | HGNC Symbol | Primer Sequence (5'-> 3') | Primer Sequence (3'-5') | Product Size (bp) |
|---|---|---|---|---|
| Signal Receptor Ligand Activity | LTA | ACCATTTCAGGGGTCGTCAC | GGATGGTTCAGGGAGGTGTG | 131 |
| Signal Receptor Ligand Activity | MIA | CAGGAGTGCAGCCACCCTAT | AAATAGCCCAGGCGAGCAG | 194 |
| Signal Receptor Ligand Activity | TNF | CAAGGACAGCAGAGGACCAG | TCCTTTCCAGGGGAGAGAGG | 156 |
| Heat Acclimation | HSPA1A | AGCTGGAGCAGGTGTGTAAC | CAGCAATCTTGGAAAGGCCC | 154 |
| Heat Acclimation | HSPA6 | CCAGAGGAACGCCACTATCC | GGAGGGATGCCACTGAGTTC | 156 |
| Heat Acclimation | BAG3 | AAGCCCAGAAGACGCACTAC | GACAGATGACCTGAACGGGG | 131 |
| Heat Acclimation | DNAJB1 | GGCTCACCTGGGCTCG | CCCGGGAATTATCCCAACCC | 129 |
| Negative Control | ACTB | CTTCGCGGGCGACGAT | CCACATAGGAATCCTTCTGACC | 104 |
| Negative Control | GAPDH | CGGGAAGGAAATGAATGGGC | GGAAAAGCATCACCCGGAGG | 148 |

**Supplementary Table 4.** List of the 13 differentially expressed genes (|log2FC| > 0.5; p-adj. < 0.05) conserved across both batches, all three cell lines, and all condition comparisons, along with descriptions of their associated biological processes.

| HGNC Symbol | Gene Name | Biological Process |
|---|---|---|
| CNTF | Ciliary Neurotrophic Factor | Promotes survival of in vitro and in vivo neuronal cell types |
| FGF18 | Fibroblast Growth Factor 18 | Involved in embryonic development, cell growth, morphogenesis, tissue repair, tumor growth, and invasion |
| GNRH1 | Gonadotropin-Releasing Hormone 1 | This gene encodes a proteolytically processed preproprotein to generate a peptide member of the gonadotropin-releasing hormone (GnRH) family of peptides. |
| GNRH2 | Gonadotropin-Releasing Hormone 2 | Codes for a preprotein, however, translation in humans has not yet been shown. |
| HBEGF | Heparin Binding EGF-like Growth Factor | Enables growth factor activity and heparin binding. It is located in the cell surface and extracellular space. |
| JAG1 | Jagged canonical notched ligand 1 | Human jagged 1 is the ligand for the receptor notch 1; the latter is involved in signaling processes. |
| LTA | Lymphotoxin Alpha | The encoded protein, a tumor necrosis factor family member, is a cytokine lymphocytes produce. The protein is highly inducible and secreted and forms heterotrimers with lymphotoxin-beta, which anchor lymphotoxin-alpha to the cell surface. This protein also mediates a large variety of inflammatory, immunostimulatory, and antiviral responses, is involved in the formation of secondary lymphoid organs during development and plays a role in apoptosis. |
| MIA | MIA SH3 domain containing | Predicted to enable growth factor activity. Predicted to be involved in extracellular matrix organization. Predicted to act upstream of or within cell-matrix adhesion. Predicted to be located in extracellular space |
| PGF | Placental Growth Factor | Enables growth factor activity. It is involved in positive regulation of cell population proliferation. It is predicted to be located in the extracellular region. Predicted to be active in extracellular space |
| PSPN | Persephin | This gene encodes a secreted ligand of the GDNF (glial cell line-derived neurotrophic factor) subfamily and TGF-beta (transforming growth factor-beta) superfamily of proteins. The encoded preproprotein is proteolytically processed to generate the mature protein. This protein signals through the RET receptor tyrosine kinase and a GPI-linked coreceptor and promotes the survival of neuronal populations. |
| SEMA4D | Semaphorin 4D | Enables identical protein binding activity, semaphorin receptor binding activity, and transmembrane signaling receptor activity. It involves several processes, including positive phosphatidylinositol 3-kinase signaling, neuron projection development regulation, and phosphate metabolic process regulation. It is an integral component of the plasma membrane. |

| | | |
|---|---|---|
| SEMA7A | Semaphorin 7A | This gene encodes a member of the semaphorin family of proteins. The encoded preproprotein is proteolytically processed to generate the mature glycosylphosphatidylinositol (GPI)-anchored membrane glycoprotein. The encoded protein is found on activated lymphocytes and erythrocytes and may be involved in immunomodulatory and neuronal processes. |
| TNF | Tumor Necrosis Factor | This gene encodes a multifunctional proinflammatory cytokine that belongs to the tumor necrosis factor (TNF) superfamily. Macrophages mainly secrete this cytokine. It can bind to and thus function through its TNFRSF1A/TNFR1 and TNFRSF1B/TNFBR receptors. This cytokine regulates a broad spectrum of biological processes, including cell proliferation, differentiation, apoptosis, lipid metabolism, and coagulation. |

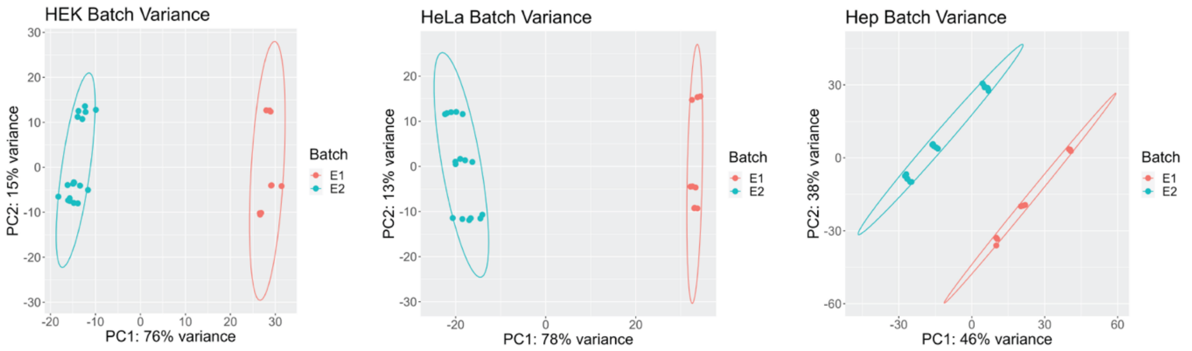


**Supplementary Figure 1. PCA Analysis Highlighting Batch Effects in Cell Lines**. PCA analyses of VST-normalized counts separated by cell line indicate that system variance is primarily driven by batch effects in HEK293 (Figure 1A), HeLa (Figure 1B), and HepG2 (Figure 1C) cells. The PCA explains 76%, 78%, and 46% of the variance in each cell line, respectively, demonstrating the impact of batch variability on gene expression profiles.

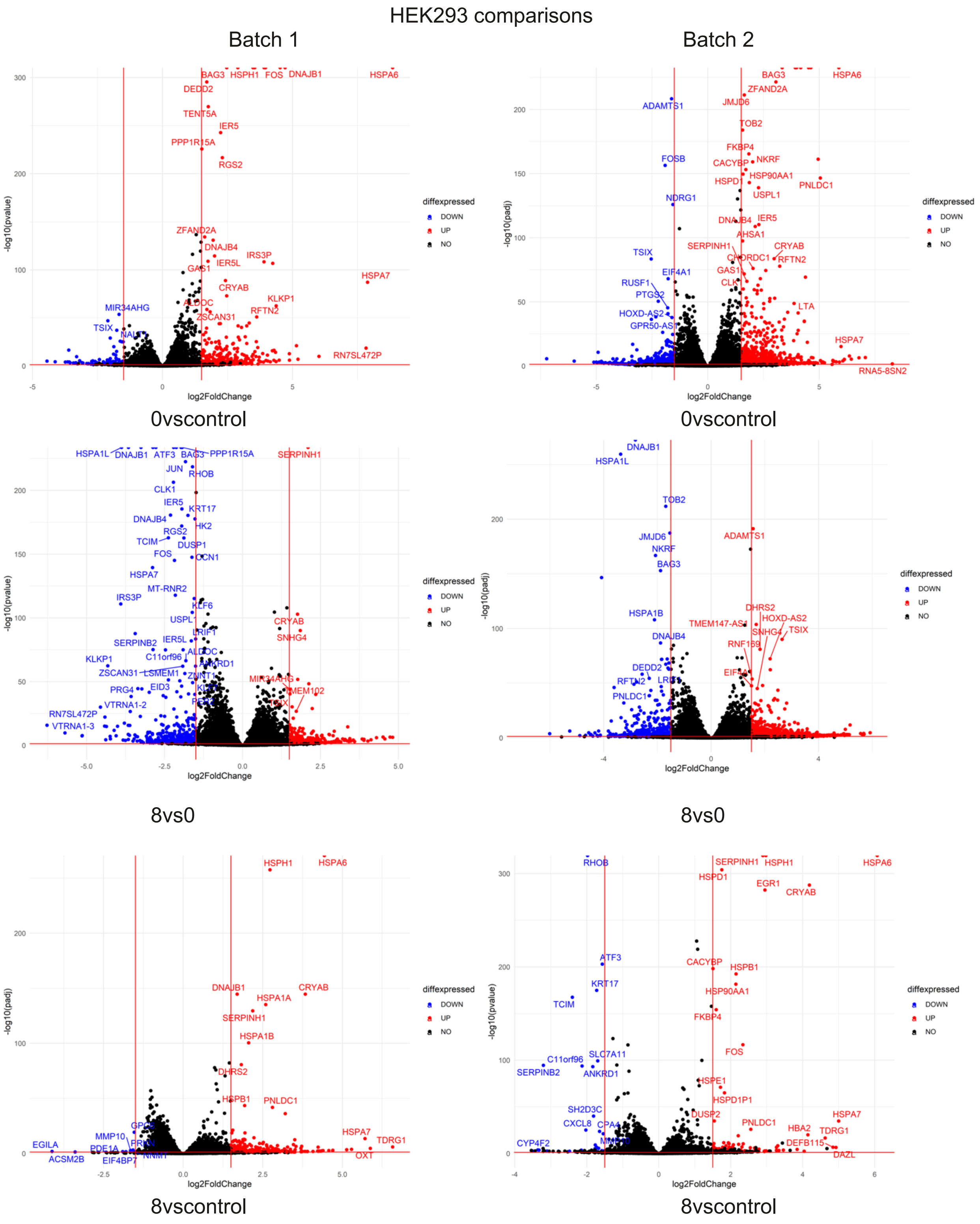
HEK293 comparisons
Batch 1
Batch 2
0vscontrol
0vscontrol
8vs0
8vs0
8vscontrol
8vscontrol
log2FoldChange
-log10(pvalue)
-log10(padj)
diffexpressed
DOWN
UP
NO

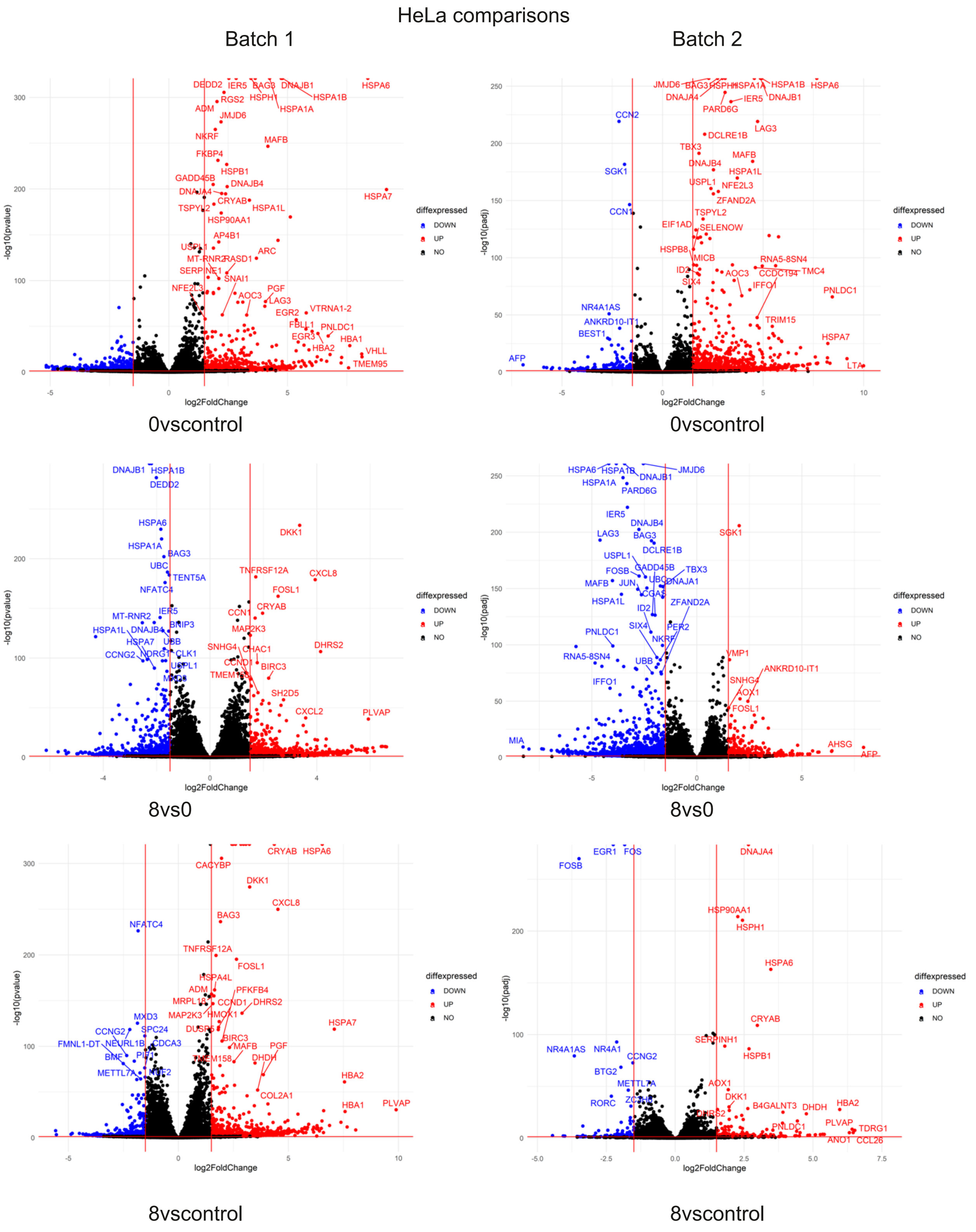
HeLa comparisons
Batch 1
Batch 2
-log10(pvalue)
-log10(padj)
log2FoldChange
diffexpressed
DOWN
UP
NO
0vscontrol
0vscontrol
8vs0
8vs0
8vscontrol
8vscontrol
DEDD2
IER5
BAG3
DNAJB1
HSPA6
HSPA1B
HSPA1A
HSPH1
RGS2
ADM
JMJD6
NKRF
MAFB
FKBP4
HSPB1
GADD45B
DNAJB4
DNAJA4
CRYAB
HSPA7
HSPA1L
TSPYL2
HSP90AA1
AP4B1
USPL1
ARC
MT-RNR2
RASD1
SNAI1
NFE2L3
PGF
AOC3
LAG3
EGR2
VTRNA1-2
PNLDC1
EGR3
HBA1
HBA2
VHLL
TMEM95
CCN2
PARD6G
DCLRE1B
TBX3
SGK1
ZFAND2A
CCN1
EIF1AD
SELENOW
HSPB8
MICB
RNA5-8SN4
TMC4
CCDC194
ID2
SIX4
IFFO1
NR4A1AS
ANKRD10-IT1
BEST1
TRIM15
AFP
LTA
DKK1
TNFRSF12A
CXCL8
UBC
TENT5A
FOSL1
NFATC4
DHRS2
CHAC1
BIRC3
SH2D5
CXCL2
PLVAP
CCNG2
CLK1
SNHG4
FOSB
JUN
CGAS
PER2
UBB
VMP1
AOX1
AHSG
CACYBP
PFKFB4
CCND1
HMOX1
DHDH
COL2A1
EGR1
FOS
HSPH1
SERPINH1
BTG2
B4GALNT3
TDRG1
CCL26
ANO1

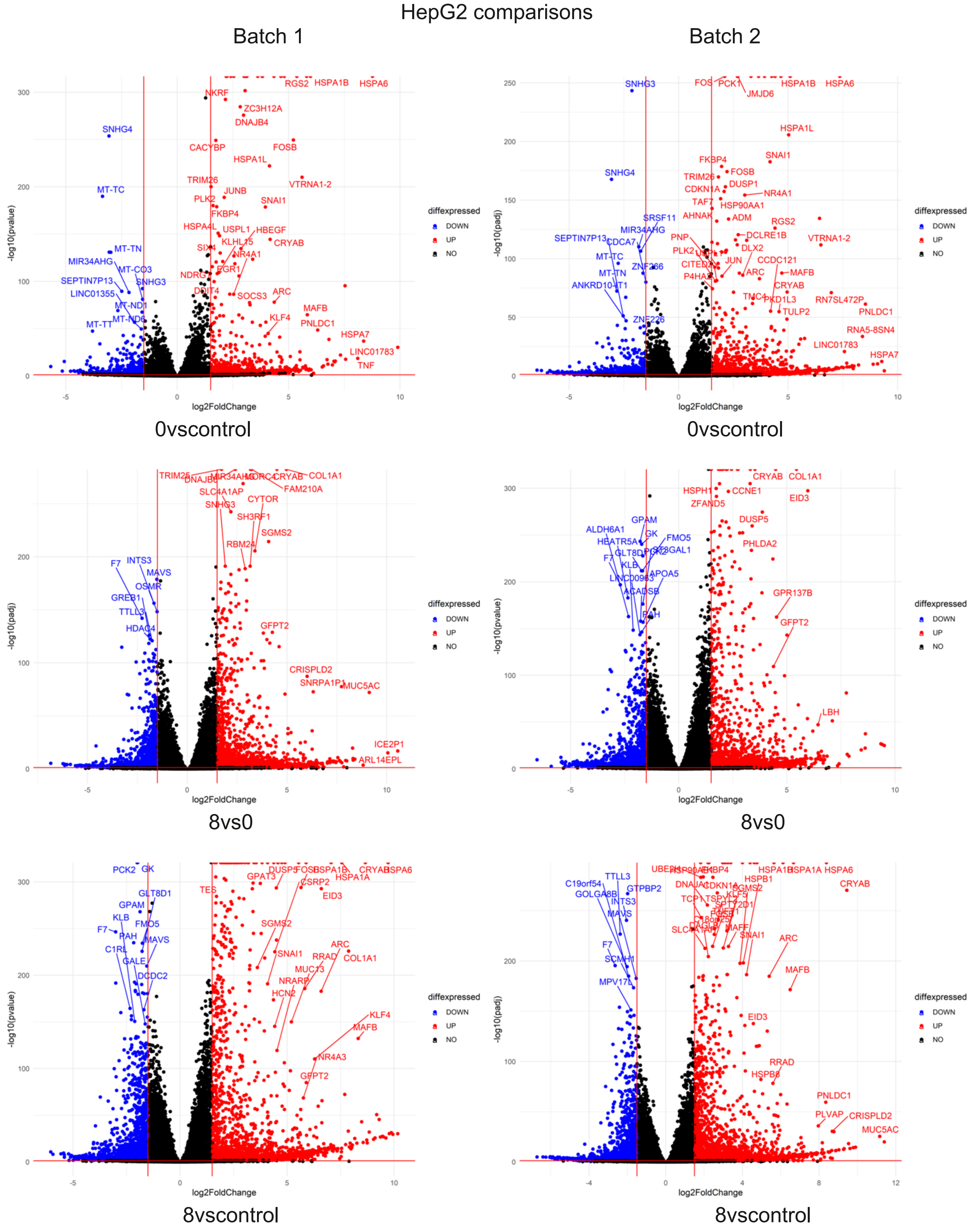


**Supplementary Figure 2. Volcano Plots of Differentially Expressed Genes by Batch**. Volcano plots illustrate the condition comparisons within each cell line for both Batch 1 and Batch 2. Each dot represents an individual gene, with blue dots indicating downregulated genes (log2 fold change < -1) and red dots indicating upregulated genes (log2 fold change > 1). Plots show gene expression changes for HEK293, HeLa, and HepG2 (all comparisons for both batches). Abbreviations are as follows: 0vsControl

(0 hours after heat shock vs. Control cells); 8vs0 (8 hours recovery after heat shock vs. 0 hours after heat shock cells); 8vsControl (8 hours recovery after heat shock vs. Control cells).

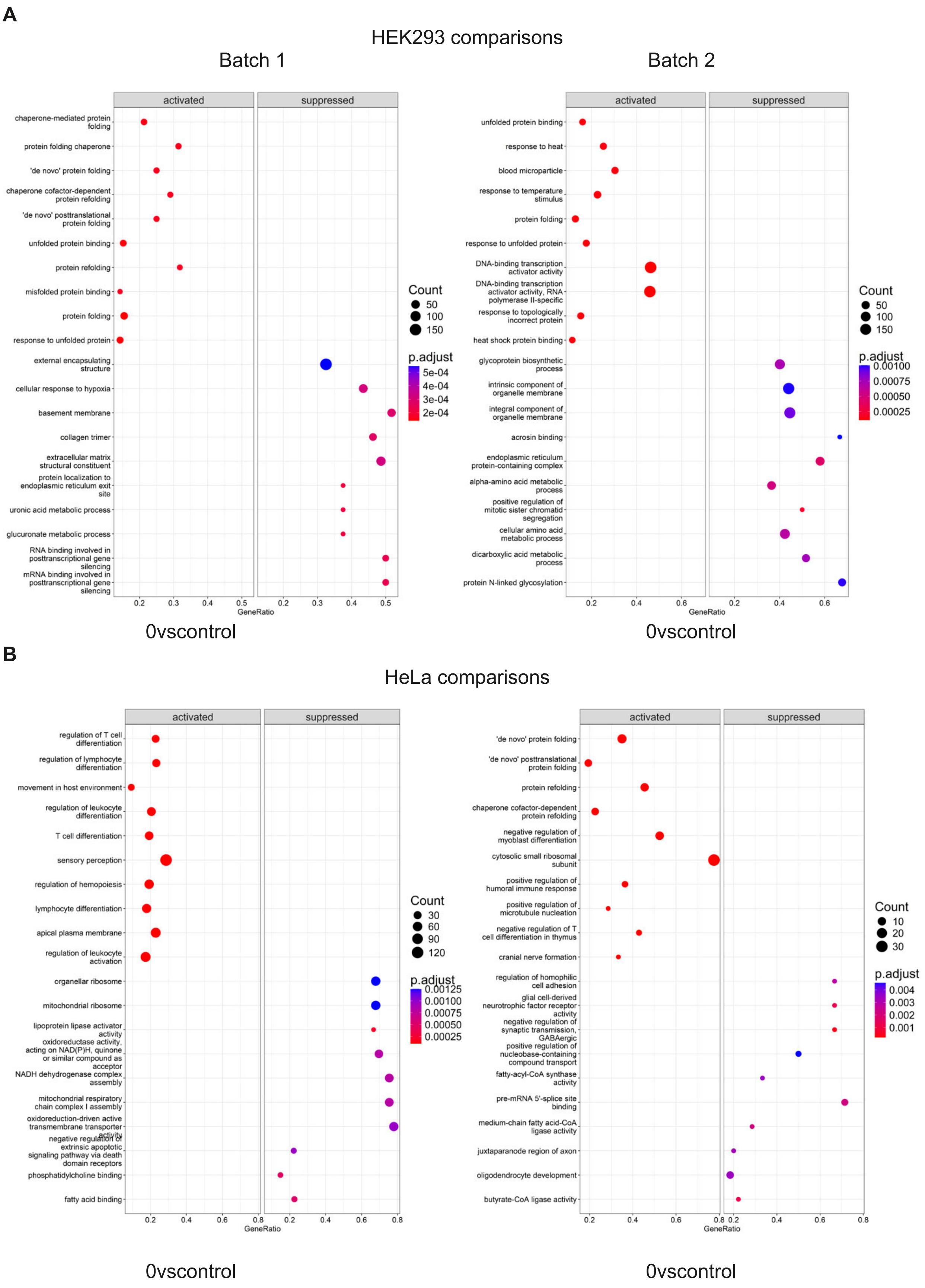

A
HEK293 comparisons
Batch 1
Batch 2
activated
suppressed
chaperone-mediated protein folding
protein folding chaperone
'de novo' protein folding
chaperone cofactor-dependent protein refolding
'de novo' posttranslational protein folding
unfolded protein binding
protein refolding
misfolded protein binding
protein folding
response to unfolded protein
external encapsulating structure
cellular response to hypoxia
basement membrane
collagen trimer
extracellular matrix structural constituent
protein localization to endoplasmic reticulum exit site
uronic acid metabolic process
glucuronate metabolic process
RNA binding involved in posttranscriptional gene silencing
mRNA binding involved in posttranscriptional gene silencing
Count
50
100
150
p.adjust
5e-04
4e-04
3e-04
2e-04
GeneRatio
0vscontrol
unfolded protein binding
response to heat
blood microparticle
response to temperature stimulus
protein folding
response to unfolded protein
DNA-binding transcription activator activity
DNA-binding transcription activator activity, RNA polymerase II-specific
response to topologically incorrect protein
heat shock protein binding
glycoprotein biosynthetic process
intrinsic component of organelle membrane
integral component of organelle membrane
acrosin binding
endoplasmic reticulum protein-containing complex
alpha-amino acid metabolic process
positive regulation of mitotic sister chromatid segregation
cellular amino acid metabolic process
dicarboxylic acid metabolic process
protein N-linked glycosylation
Count
50
100
150
p.adjust
0.00100
0.00075
0.00050
0.00025
GeneRatio
0vscontrol
B
HeLa comparisons
activated
suppressed
regulation of T cell differentiation
regulation of lymphocyte differentiation
movement in host environment
regulation of leukocyte differentiation
T cell differentiation
sensory perception
regulation of hemopoiesis
lymphocyte differentiation
apical plasma membrane
regulation of leukocyte activation
organellar ribosome
mitochondrial ribosome
lipoprotein lipase activator activity
oxidoreductase activity, acting on NAD(P)H, quinone or similar compound as acceptor
NADH dehydrogenase complex assembly
mitochondrial respiratory chain complex I assembly
oxidoreduction-driven active transmembrane transporter activity
negative regulation of extrinsic apoptotic signaling pathway via death domain receptors
phosphatidylcholine binding
fatty acid binding
Count
30
60
90
120
p.adjust
0.00125
0.00100
0.00075
0.00050
0.00025
GeneRatio
0vscontrol
'de novo' protein folding
'de novo' posttranslational protein folding
protein refolding
chaperone cofactor-dependent protein refolding
negative regulation of myoblast differentiation
cytosolic small ribosomal subunit
positive regulation of humoral immune response
positive regulation of microtubule nucleation
negative regulation of T cell differentiation in thymus
cranial nerve formation
regulation of homophilic cell adhesion
glial cell-derived neurotrophic factor receptor activity
negative regulation of synaptic transmission, GABAergic
positive regulation of nucleobase-containing compound transport
fatty-acyl-CoA synthase activity
pre-mRNA 5'-splice site binding
medium-chain fatty acid-CoA ligase activity
juxtaparanode region of axon
oligodendrocyte development
butyrate-CoA ligase activity
Count
10
20
30
p.adjust
0.004
0.003
0.002
0.001
GeneRatio
0vscontrol

**C**

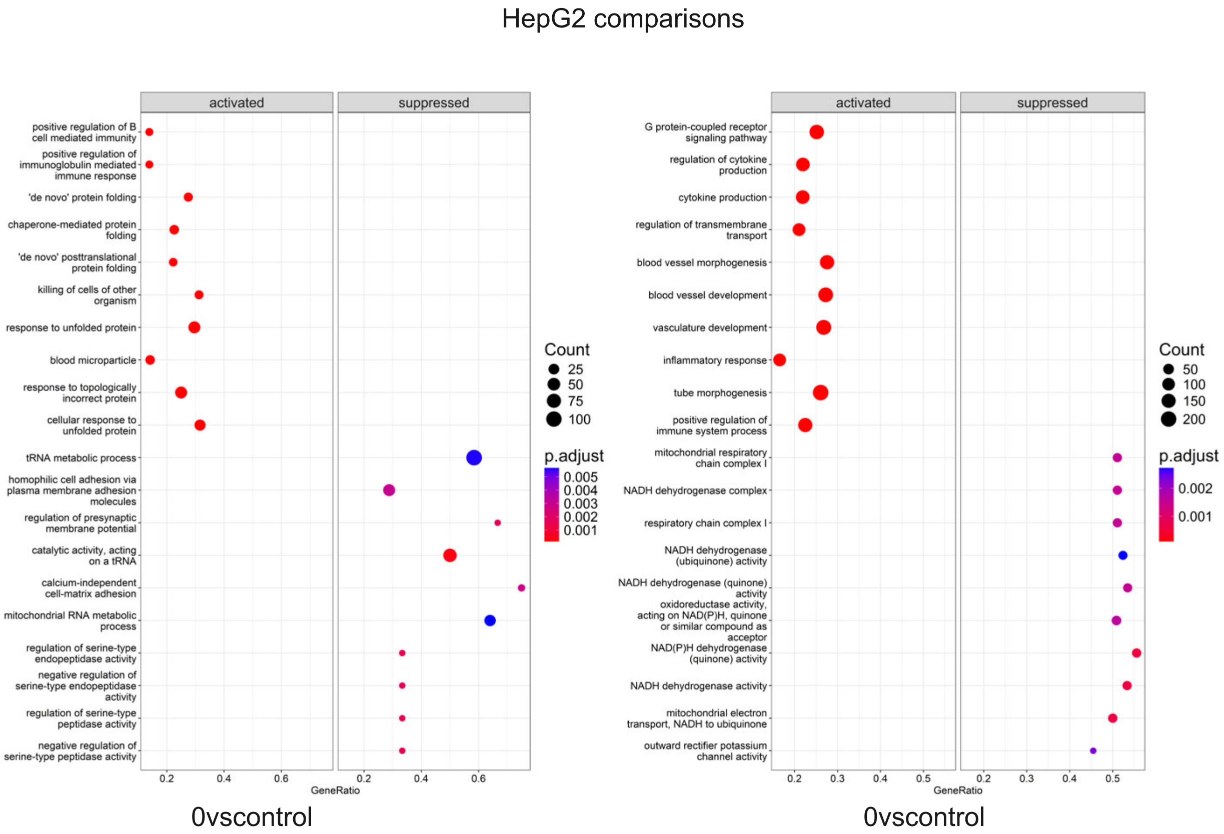
HepG2 comparisons
activated
suppressed
positive regulation of B cell mediated immunity
positive regulation of immunoglobulin mediated immune response
'de novo' protein folding
chaperone-mediated protein folding
'de novo' posttranslational protein folding
killing of cells of other organism
response to unfolded protein
blood microparticle
response to topologically incorrect protein
cellular response to unfolded protein
tRNA metabolic process
homophilic cell adhesion via plasma membrane adhesion molecules
regulation of presynaptic membrane potential
catalytic activity, acting on a tRNA
calcium-independent cell-matrix adhesion
mitochondrial RNA metabolic process
regulation of serine-type endopeptidase activity
negative regulation of serine-type endopeptidase activity
regulation of serine-type peptidase activity
negative regulation of serine-type peptidase activity
0.2
0.4
0.6
GeneRatio
Count
25
50
75
100
p.adjust
0.005
0.004
0.003
0.002
0.001
0vscontrol
activated
suppressed
G protein-coupled receptor signaling pathway
regulation of cytokine production
cytokine production
regulation of transmembrane transport
blood vessel morphogenesis
blood vessel development
vasculature development
inflammatory response
tube morphogenesis
positive regulation of immune system process
mitochondrial respiratory chain complex I
NADH dehydrogenase complex
respiratory chain complex I
NADH dehydrogenase (ubiquinone) activity
NADH dehydrogenase (quinone) activity
oxidoreductase activity, acting on NAD(P)H, quinone or similar compound as acceptor
NAD(P)H dehydrogenase (quinone) activity
NADH dehydrogenase activity
mitochondrial electron transport, NADH to ubiquinone
outward rectifier potassium channel activity
0.2
0.3
0.4
0.5
GeneRatio
Count
50
100
150
200
p.adjust
0.002
0.001
0vscontrol

D

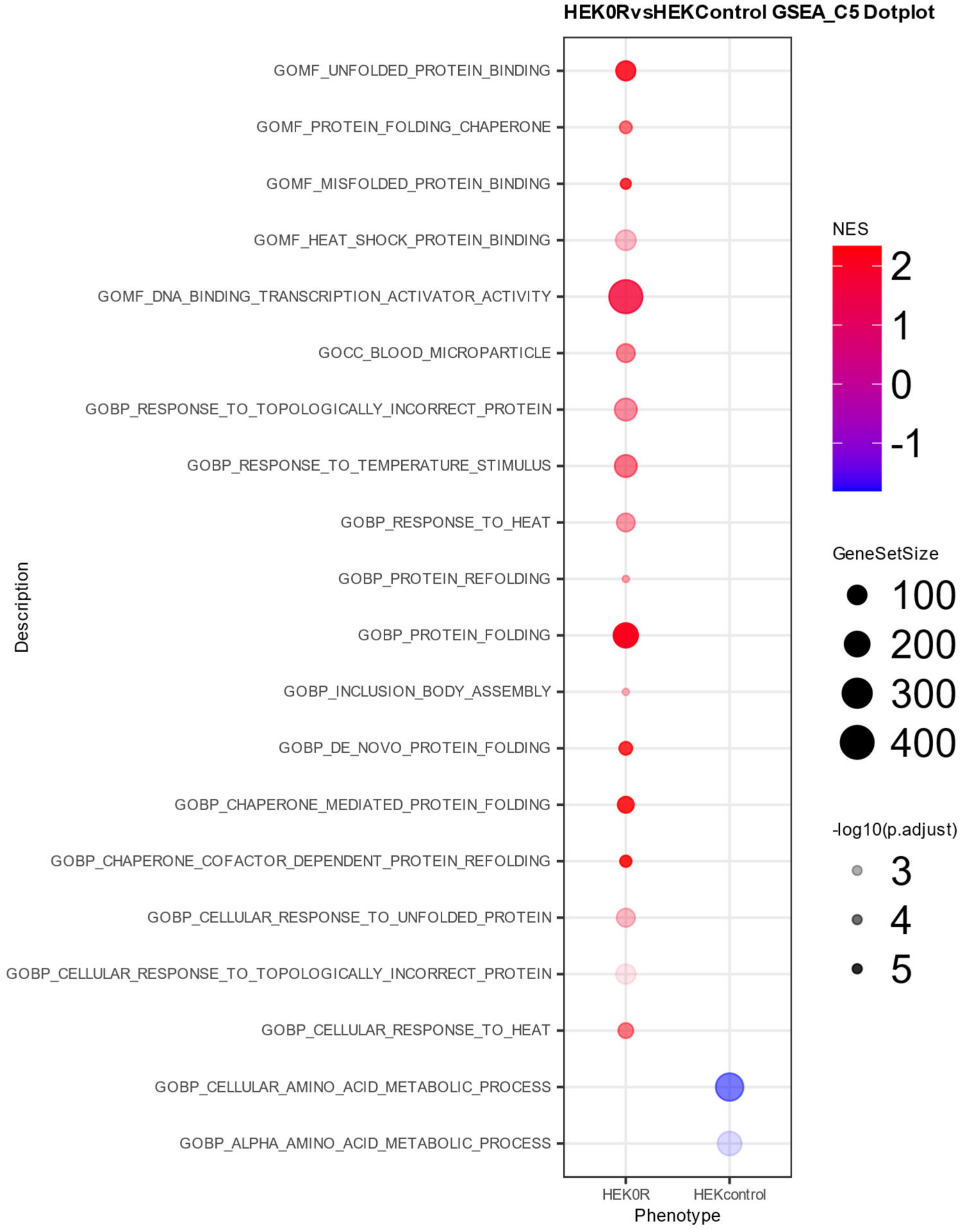
HEK0RvsHEKControl GSEA_C5 Dotplot
GOMF_UNFOLDED_PROTEIN_BINDING
GOMF_PROTEIN_FOLDING_CHAPERONE
GOMF_MISFOLDED_PROTEIN_BINDING
GOMF_HEAT_SHOCK_PROTEIN_BINDING
GOMF_DNA_BINDING_TRANSCRIPTION_ACTIVATOR_ACTIVITY
GOCC_BLOOD_MICROPARTICLE
GOBP_RESPONSE_TO_TOPOLOGICALLY_INCORRECT_PROTEIN
GOBP_RESPONSE_TO_TEMPERATURE_STIMULUS
GOBP_RESPONSE_TO_HEAT
GOBP_PROTEIN_REFOLDING
GOBP_PROTEIN_FOLDING
GOBP_INCLUSION_BODY_ASSEMBLY
GOBP_DE_NOVO_PROTEIN_FOLDING
GOBP_CHAPERONE_MEDIATED_PROTEIN_FOLDING
GOBP_CHAPERONE_COFACTOR_DEPENDENT_PROTEIN_REFOLDING
GOBP_CELLULAR_RESPONSE_TO_UNFOLDED_PROTEIN
GOBP_CELLULAR_RESPONSE_TO_TOPOLOGICALLY_INCORRECT_PROTEIN
GOBP_CELLULAR_RESPONSE_TO_HEAT
GOBP_CELLULAR_AMINO_ACID_METABOLIC_PROCESS
GOBP_ALPHA_AMINO_ACID_METABOLIC_PROCESS
Description
HEK0R
HEKcontrol
Phenotype
NES
2
1
0
-1
GeneSetSize
100
200
300
400
-log10(p.adjust)
3
4
5

**E**

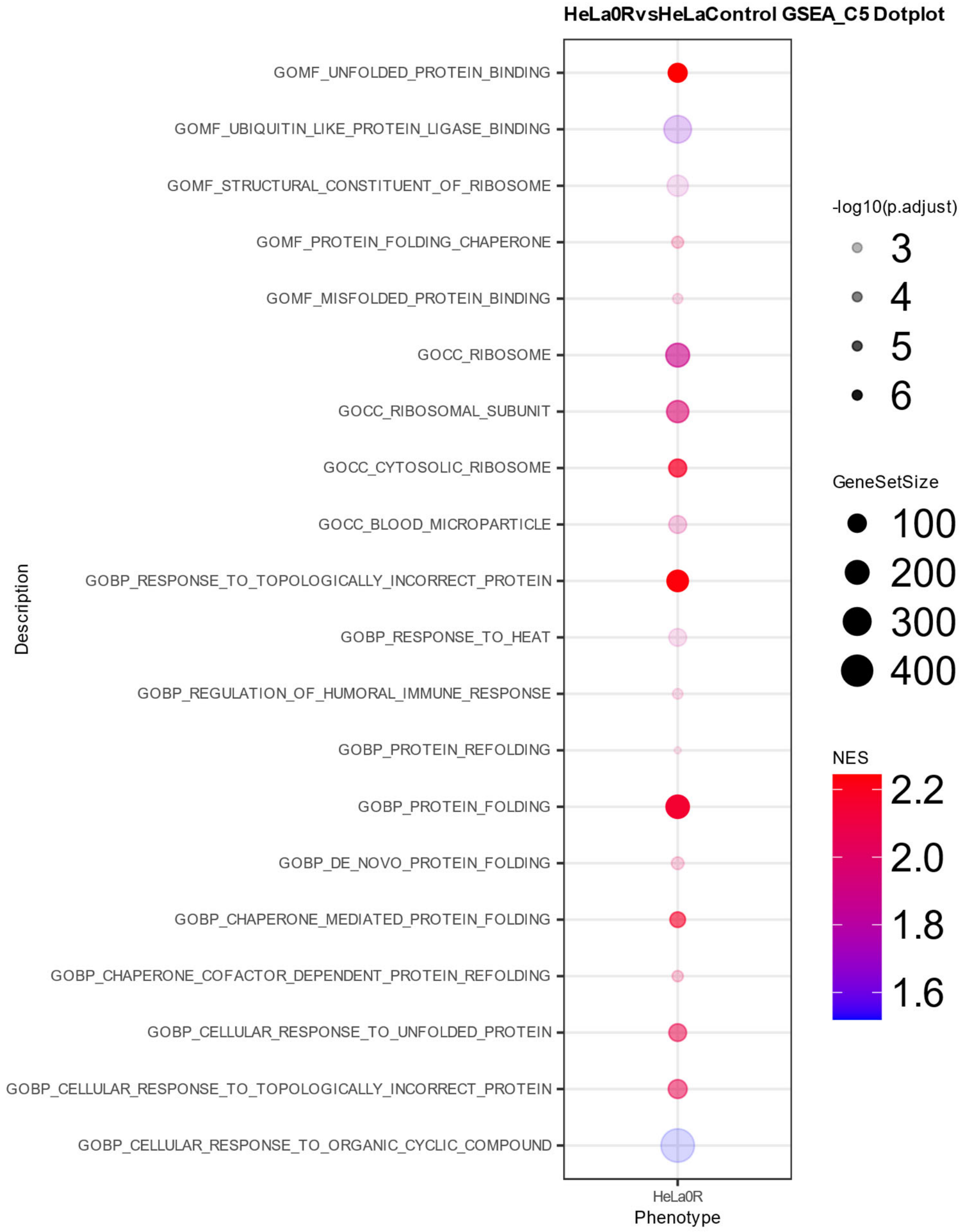
HeLa0RvsHeLaControl GSEA_C5 Dotplot
GOMF_UNFOLDED_PROTEIN_BINDING
GOMF_UBIQUITIN_LIKE_PROTEIN_LIGASE_BINDING
GOMF_STRUCTURAL_CONSTITUENT_OF_RIBOSOME
GOMF_PROTEIN_FOLDING_CHAPERONE
GOMF_MISFOLDED_PROTEIN_BINDING
GOCC_RIBOSOME
GOCC_RIBOSOMAL_SUBUNIT
GOCC_CYTOSOLIC_RIBOSOME
GOCC_BLOOD_MICROPARTICLE
GOBP_RESPONSE_TO_TOPOLOGICALLY_INCORRECT_PROTEIN
GOBP_RESPONSE_TO_HEAT
GOBP_REGULATION_OF_HUMORAL_IMMUNE_RESPONSE
GOBP_PROTEIN_REFOLDING
GOBP_PROTEIN_FOLDING
GOBP_DE_NOVO_PROTEIN_FOLDING
GOBP_CHAPERONE_MEDIATED_PROTEIN_FOLDING
GOBP_CHAPERONE_COFACTOR_DEPENDENT_PROTEIN_REFOLDING
GOBP_CELLULAR_RESPONSE_TO_UNFOLDED_PROTEIN
GOBP_CELLULAR_RESPONSE_TO_TOPOLOGICALLY_INCORRECT_PROTEIN
GOBP_CELLULAR_RESPONSE_TO_ORGANIC_CYCLIC_COMPOUND
Description
HeLa0R
Phenotype
-log10(p.adjust)
3
4
5
6
GeneSetSize
100
200
300
400
NES
2.2
2.0
1.8
1.6

**F**

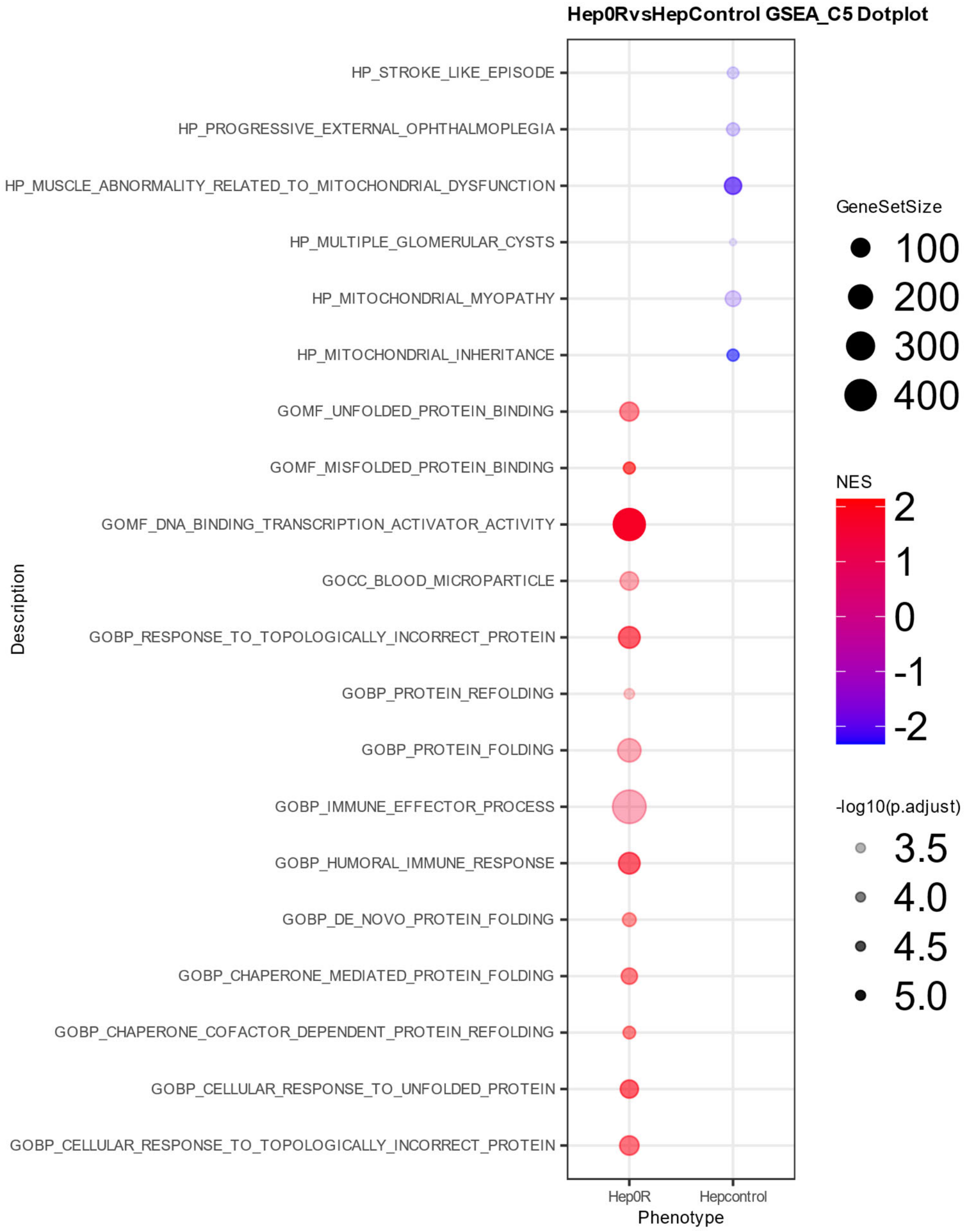


**Supplementary Figure 3. Gene Set Enrichment Analysis via Dot Plots**. Dot plots visualize Gene Set Enrichment Analysis (GSEA) results for the top 15 positive and top 15 negative normalized enrichment score (NES) hits [using the Human Phenotype Ontology (A-C) and the geneOntology databases (GO) (D-F)] in HEK293 (A, D), HeLa (B, E), and HepG2 (C, F) cell lines at 0 hours vs control cells, providing insights into enriched pathways influenced by heat shock. Abbreviations are 0vsControl (0 hours after heat shock vs. Control cells).

HEK293 comparisons

Batch 1

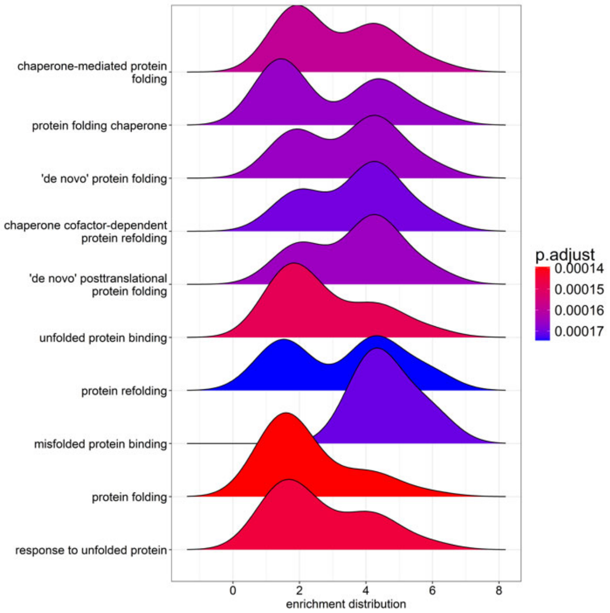


0vscontrol

Batch 2

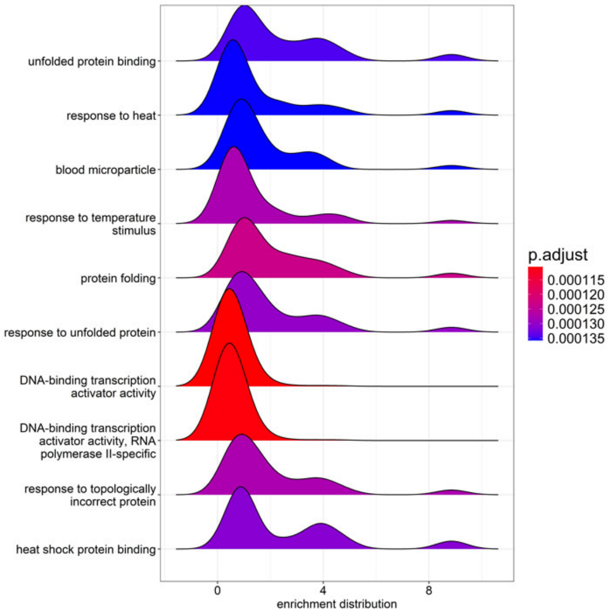


0vscontrol

HeLa comparisons

Batch 1

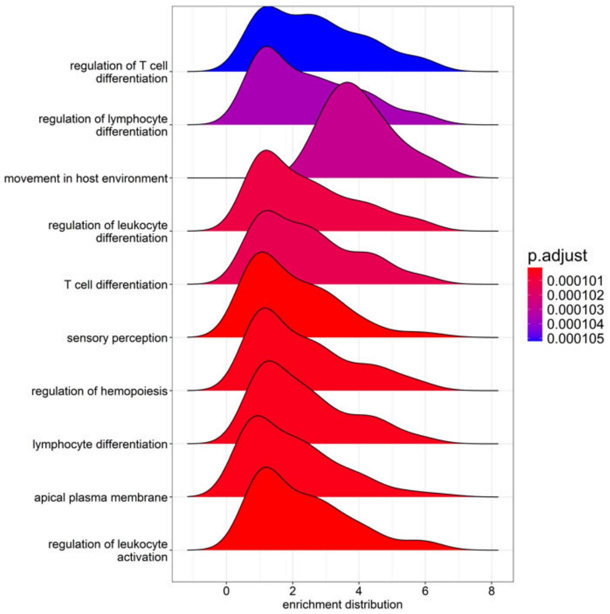


0vscontrol

Batch 2

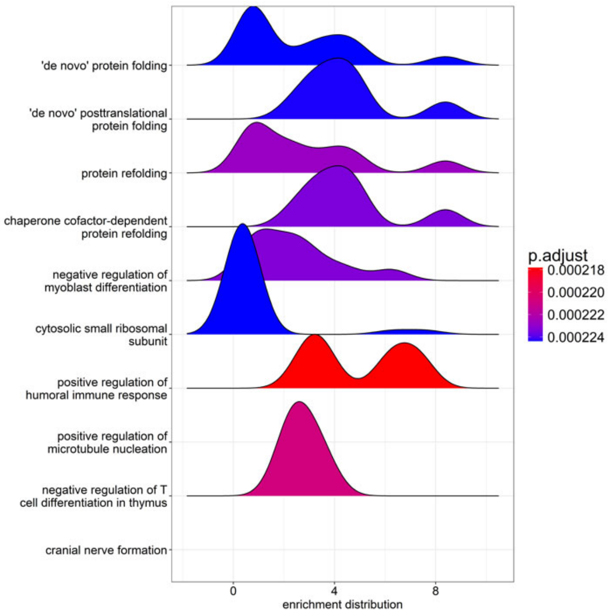


0vscontrol

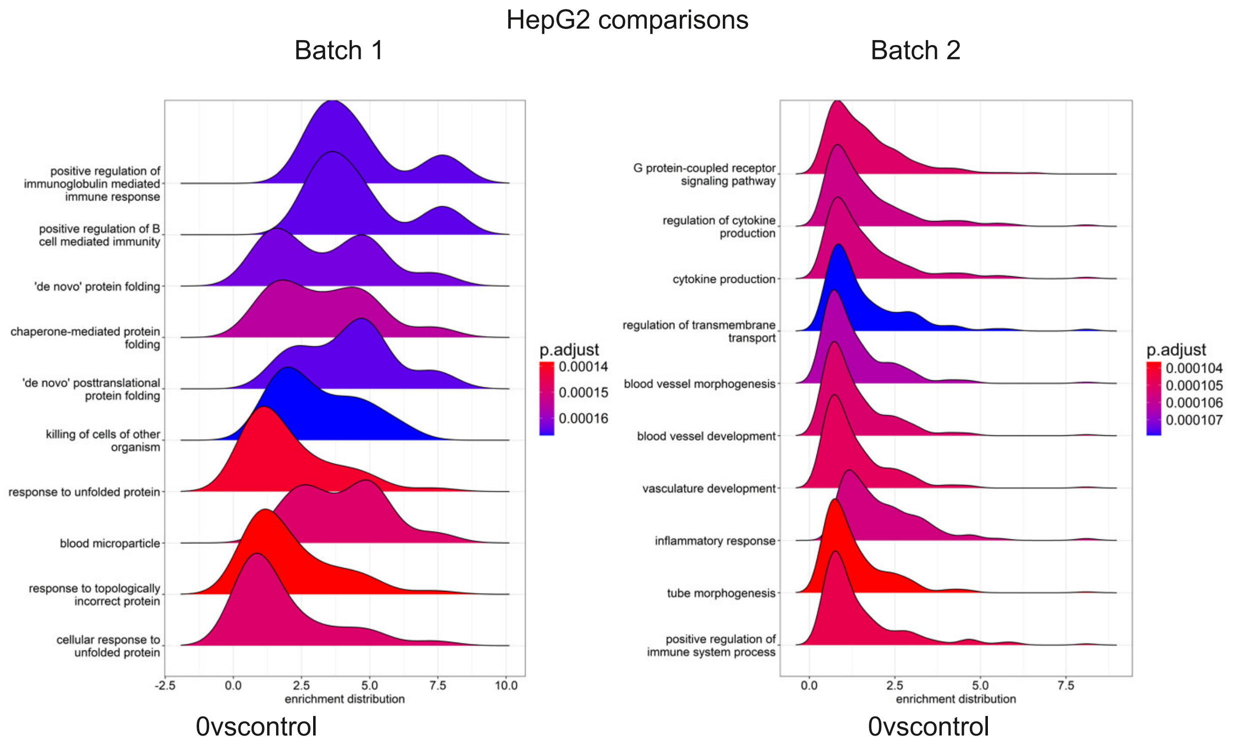


**Supplementary Figure 4. Distribution of Enriched Genes Across Top GSEA Hits**. Distribution plots of log2 fold changes for genes in the top 15 positively enriched GSEA pathways for HEK293, HeLa, and HepG2 cell lines at 0 hours vs. Control. Peak height indicates the number of enriched genes within each log2 fold change range. Abbreviations are as follows: 0vsControl (0 hours after heat shock vs. Control cells).

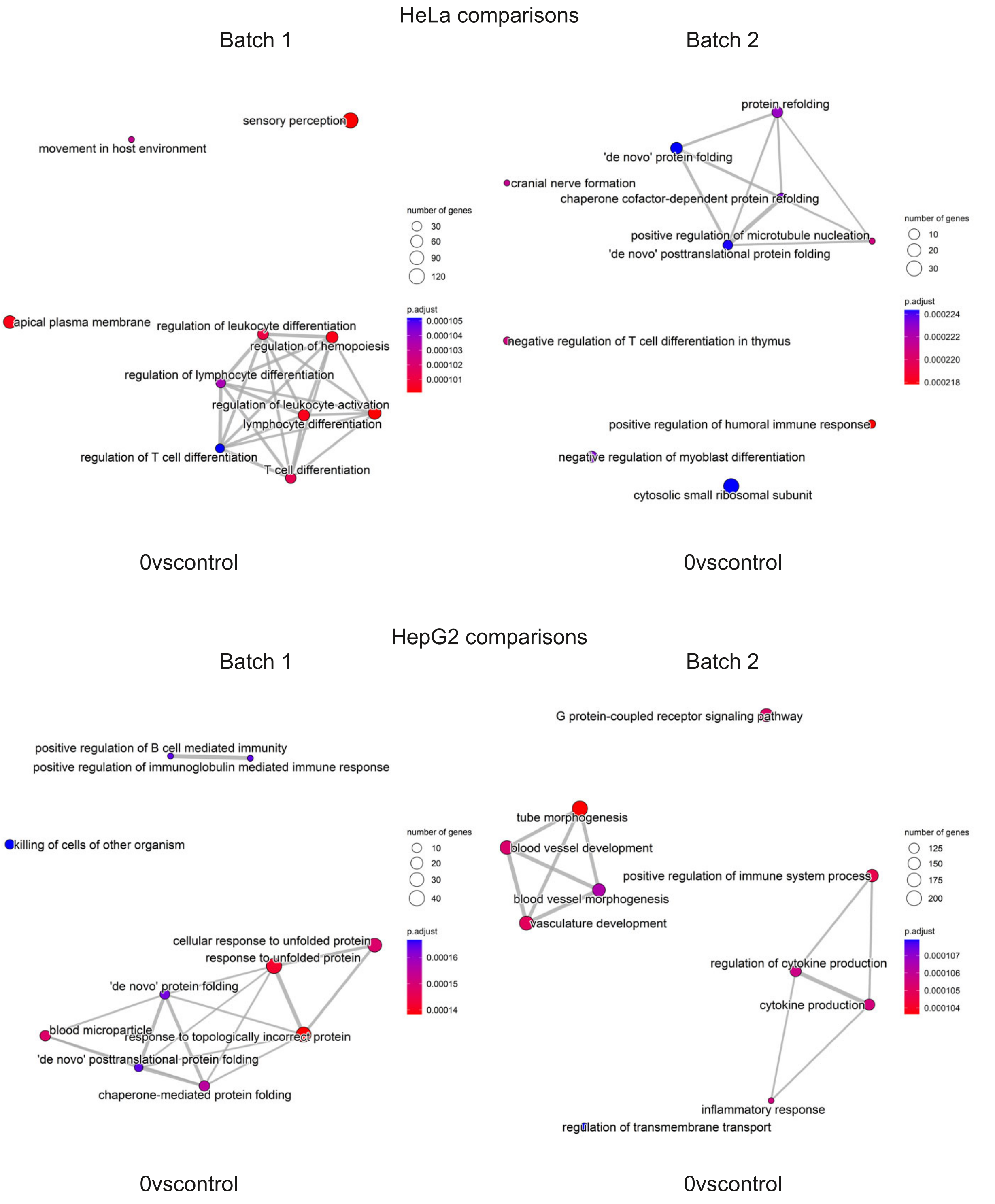


**Supplementary Figure 5. Enrichment Maps of GSEA Hits for HeLa and HepG2 Cells**. Network enrichment maps of the top 10 positive GSEA NES hits for HeLa and HepG2 cell lines at 0 hours vs. Control. Nodes represent enriched gene sets, while edges indicate shared genes between sets. Maps are shown for HeLa (Batch 1, top left panel; Batch 2, top right panel) and HepG2 (Batch 1, bottom left panel; Batch 2, bottom right panel), providing a systems-level perspective on pathway relationships under heat shock conditions.

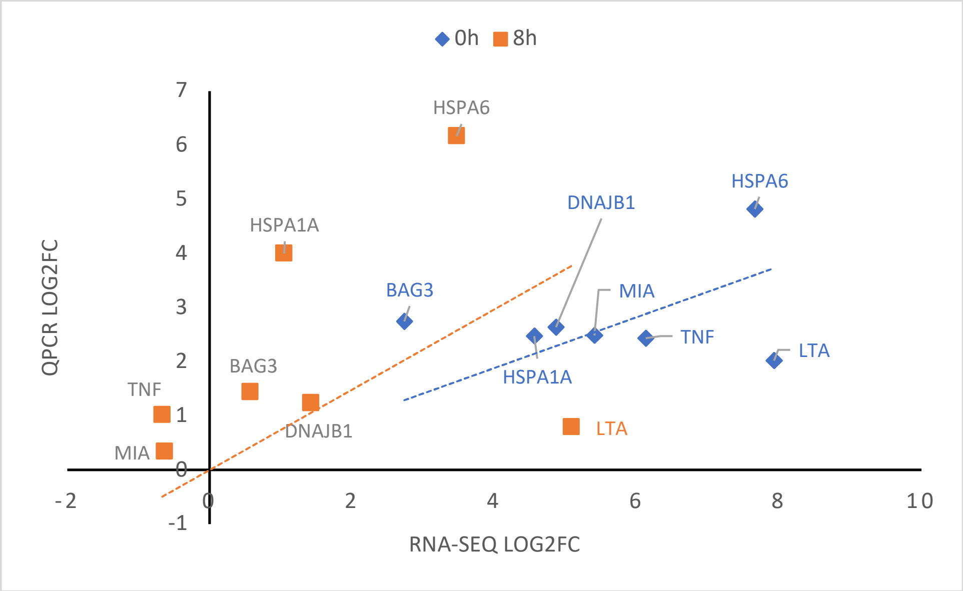


**Supplementary Figure 6. Scatter plot comparing RNA-seq and qPCR fold change values.** The scatter plot displays log2 fold change (log2FC) values obtained from RNA-seq (x-axis) and qPCR (y-axis) analyses for genes HSPA1A, HSPA6, BAG3, DNAJB1, LTA, MIA, and TNF at 0 hours (blue circles) and 8 hours (orange squares) post-heat shock. Data points near the diagonal indicate strong agreement between RNA-seq and qPCR measurements, while points further from the diagonal represent differences in the magnitude of fold change between the two methods, suggesting potential differences in sensitivity or dynamic range.
