## supplementa methods for "Transcriptomics Unveil Canonical and Non-Canonical Heat Shock-Induced Pathways in Human Cell Lines"

**Detailed Transcriptomics Analysis**

**Directory Organization and Tool Installation**

Paired sequencing reads, provided as compressed FASTQ.gz files by Novogene (as detailed in "Sample Preparation, cDNA Library Preparation, and Sequencing"), was downloaded onto a Linux-based system. Organizational directories were created to store raw data separately, cleaned reads, the indexed genome, aligned reads, and gene counts. Software tools were installed following the instructions in each tool’s documentation.

Anaconda installed Python and Jupyter Notebook for Windows-based analysis, providing a shareable, interactive development environment for custom Python scripts. Additionally, R version 4.1.1 (2021-08-10) "Kick Things" and R Studio (2022.07.1 Build 554 "Spotted Wakerobin" Release, 7872775e) were installed for running R scripts within a dedicated project and environment.

**Read Quality Control Overview**

Adapter sequences, PhiX library sequences, and low-quality reads are common byproducts of library preparation and next-generation sequencing (NGS). To ensure that high-quality reads are used for downstream analysis, BBDuk was employed to perform adapter trimming, PhiX filtering, and quality filtering based on Phred scores [43,44]. This process took paired compressed FASTQ.gz files as input and, via command-line execution, removed any artifacts from sequencing or library preparation to prevent them from being included in the expression data for further analysis.

**Adapter Trimming**

To eliminate sequenced adapter reads from downstream processing and analysis, we used BBDuk’s k-mer trimming functionality. Each paired read was processed using a BBDuk shell script, which removed reads that matched a reference k-mer corresponding to an adapter sequence. The resulting cleaned FASTQ.gz files were saved back into the raw data directory, with filenames appended as “(pair 1 or pair 2)clean.fq.gz.” The outm option was used to store filtered adapter sequences, and the ref option specified the file containing adapter sequences to be filtered. The trimming was set to right-side (ktrim=r), with a k-mer size of 23, a minimum k-mer size (mink) of 11, and a hamming distance (hdist) of 1. The tbo flag enabled pair overlap detection using BBMerge, and the tpe flag ensured paired reads were trimmed to the same length. Summary statistics were output to a .txt file via the stats option.

**Phred Quality Score and PhiX Trimming**

BBDuk was utilized to eliminate reads that matched the k-mers associated with the PhiX control library, like the adapter trimming process. To ensure that only high-quality reads were included in the analysis, BBDuk’s Phred quality trimming function was applied. This function scans the right end of the paired reads and trims them until the Phred quality score meets the specified threshold; if the score remains unsatisfactory, the entire paired read is discarded. The adapter-trimmed paired reads were generated as outlined in "Read Quality Control-Adapter Trimming."

For filtering PhiX sequences and performing quality trimming, the BBDuk script was executed via command line for each pair of reads. The cleaned FASTQ.gz files were then stored in a results folder, with filenames marked as "(pair 1 or pair 2)tf.fq.gz." The outm option redirected reads that matched the PhiX k-mers to a designated file, while the ref option referenced the Illumina PhiX spike-in sequence. The k-mer size was set to 31, with a hamming distance (hdist) of 1. Additionally, summary statistics were generated and saved in a .txt file using the stats option. Quality trimming was restricted to the right side (qtrim=r), with the threshold for trimming set to Q10 (trimq=10) based on Phred scores.

**Reference Genome Indexing and Alignment of Reads to an Indexed Reference Genome**

To quantify gene expression, it is essential to map high-quality reads to the human reference genome to identify the features in the dataset. STAR was employed for this purpose, accepting reads cleaned by BBDuk and outputting binary sequence alignment map files (.BAM) after performing splice-aware mapping against the indexed human reference genome (GRCh38.p13) [45]. Before alignment, the indexed reference genome was generated.

The STAR command was executed from the command line, with the runThreadN parameter set to 24 and the runMode set to “genomeGenerate.” The genomeDir specified the directory for the indexed genome, while genomeFastaFiles pointed to the folder containing the GRCh38.p13 reference genome in FASTA format. The sjdbGTFfile was set to reference the Gencode primary assembly annotation (.gtf file), and sjdbOverhang was configured to “99.” This indexed reference genome was created once and subsequently used for all alignments.

Compressed, paired, and adapter-trimmed reads that had undergone PhiX and quality trimming were produced as outlined in “Adapter trimming” and “Phred Quality Score and PhiX Trimming.” For two-pass mapping, STAR was again executed via the command line, with runThreadN set to “24” and runMode specified as “alignReads.” The twopassMode was configured to “Basic,” while genomeDir indicated the directory containing the previously generated indexed reference genome. The readFilesIn parameter was set to the paths of each paired BBDuk output file (1 or 2). The genomeLoad option was set to “NOSharedMemory,” and outFileNamePrefix was formatted as CellLinesamplenumber_condition. The outSAMtype was defined as “BAM SortedByCoordinate,” and the outReadsUnmapped option was set to “Fastx” to save unmapped reads in a FASTQ file. Finally, the quantMode was configured to “GeneCounts” for secondary feature counting, and readFilesCommand was set to “zcat” to facilitate reading compressed .gz files.

**Sort and Index Mapped Reads for Feature Counting**

Samtools was used to sort an input .BAM file and create a corresponding index file (.BAI), which is necessary for downstream feature counting steps [46]. The splice-aware and mapped .BAM files generated by STAR, as detailed in “Reference Genome Indexing and Alignment of Reads to an Indexed Reference Genome,” served as the input.

To sort the .BAM file, the Samtools sort command was executed from the command line in the same directory as the STAR output files. The -m flag was set to 450,000,000, the -o flag specified the output format as CellLinesamplenumberconditionAligned.out.bam, and the -O flag indicated that the output should be in BAM format. The thread count was set to 24 using the -@ flag, followed by the path to the input file.

After sorting, the Samtools index command was run from the same directory as the sorted .BAM files. The -b flag was included to generate the index, and the path to the sorted .BAM files was provided. The index was named by appending “.bai” to the name of the input file.

**Feature Counts**

HTSeq was utilized to create a raw gene count matrix by determining the number of reads mapped to each feature. The inputs required for HTSeq include a sorted aligned .BAM file, its corresponding index .BAI file, and a reference human genome annotation file (GRCh38.104.gtf) [47]. The sorted and indexed alignment files were produced as outlined in “Sort and Index Mapped Reads for Feature Counting.”

The HTSeq script was executed from the command line in the same directory where the sorted .BAM and .BAI files were located. The order parameter was set to “name,” specifying the expected format of the sorted input. The stranded option was set to “reverse,” meaning that each paired read must align to the opposite strand of the feature to properly account for the sequencing of either the template or coding strands. The feature_type was designated as “exon,” while the id_attribute was specified as “gene_id,” and additional attributes were set to “gene_name.” The mode was defined as “union,” the output counts were saved as a .csv file in a results folder. The paths to the input file and the gencode.v38.primary_assembly.annotation.gtf file were provided.

**Quality Control**

BBDuk, STAR, Samtools, and HTSeq generated multiple output files distributed across various directories. To summarize and evaluate these files, MultiQC was utilized, which provides an overview of file types associated with next-generation sequencing read processing [48]. By running MultiQC from a directory containing all relevant subdirectories with output files, users can verify the expected number of files and their types for each sample.

MultiQC was executed using the command line entry “multiqc .” within the directory containing all previously generated output files. For each sample, it presented general statistics, HTSeq-specific metrics, BBTools statistics, STAR-specific metrics, and STAR gene count statistics.

General statistics included the percentage of mapped reads and the total mapped reads in millions. HTSeq-specific metrics provided a breakdown of mapped reads by percentage, total mapped reads in millions, and detailed read classifications (assigned, ambiguous, not uniquely aligned, no feature, low quality, and not aligned). BBTools statistics reported the proportions of reads filtered based on matching k-mers. STAR-specific metrics included the counts of uniquely mapped reads, reads mapped to multiple loci, reads mapped to too many loci, and unmapped reads categorized as short reads or classified as “other.”

STAR gene counts statistics detailed the number of overlapping genes, no feature calls, ambiguous feature calls, multimapped reads, and unmapped reads, all available as percentages or in millions. These statistics were provided for unstranded, same stranded, and reverse stranded workflows.

**Generating and Visualizing Differentially Expressed Genes**

The negative binomial distribution is a suitable model for analyzing raw count data that is well-dispersed and influenced by multiple sources of variance, making DESeq2 an ideal choice for identifying differentially expressed genes (DEGs) [49]. DESeq2 estimates the mean and models the dispersion of gene expression for each gene observed in the dataset.

Once the dispersion for gene expression is determined, the counts for each gene are fitted to a negative binomial generalized linear model. In simpler terms, the coefficients obtained from this modeling represent the log fold changes in expression between specific comparisons [49]. The statistical significance of these changes in expression is assessed through a pairwise comparison method known as the Wald test, where the null hypothesis posits that the log fold change is zero.

Raw gene read counts were produced as detailed in “Feature Counts” and then imported into R. For each gene across the experimental comparisons of interest (R0 vs. Control, R8 vs. Control, R8 vs. R0), DESeq2 took the HTSeq-measured raw read counts as input and computed the differential expression between the conditions.

After loading all the required libraries, the dds object was created using the DESeq2 function DESeqDataSetFromMatrix, where countData was assigned the raw gene counts matrix, colData was linked to a table outlining the experimental design, and the design formula was set to “~ condition” to indicate the analysis axis. Levels were established within the dds object to facilitate subsequent comparison commands. Counts below 250 were filtered out, and the DESeq function was applied to the dds object.

The results function was then executed on the dds object, with alpha set to 0.05 and contrast matrices defined for the comparisons of interest (R0 vs. Control, R8 vs. Control, R8 vs. R0). The outputs were saved into new variables and exported to .csv files that included the base mean, log2 fold change (LFC), the standard error of the LFC, the Wald statistic (calculated as LFC divided by the standard error of the LFC), p-values, and adjusted p-values (p.adj) for multiple comparisons. For each comparison, the ggplot2 library from the tidyverse was used to create volcano plots displaying gene LFC against -log10(gene p.adj).

**Gene Counts Normalization and Dimensionality Reduction**

Utilizing DESeq2 for differential gene expression (DGE) analysis allows researchers to leverage its built-in variance stabilizing transformation (VST), which helps reduce the impact of outliers and addresses heteroscedasticity [49]. The VST-normalized counts were subsequently used as inputs for dimensionality reduction analyses.

Principal component analysis (PCA) and heatmap analysis were employed as dimensionality reduction techniques. These methods were chosen to effectively visualize the intricate VST-normalized gene expression data, encompassing over 20,000 gene features across nine samples, into a more concise and interpretable format. PCA was conducted to identify the primary sources of variation within the dataset and to estimate the effect sizes between samples based on standard deviation.

**Dimensionality Reduction-Principal Component Analysis**

To distill the gene features per sample into a representation that highlights the primary contributors to variance within the dataset, principal component analysis (PCA) was conducted using R’s prcomp function, along with the tidyverse and ggfortify libraries.

First, VST counts were calculated and organized into a dataframe, from which non-finite rows were removed, resulting in a clean data matrix. This data matrix was then used as input for the prcomp function, with the parameter “scale.” set to “TRUE” to ensure proper scaling. Subsequently, a PCA results object was generated. The autoplot function was employed to visualize the score plot of the first two principal components, and the plot was saved as an image file.

**Histograms, Volcano Plots, and Venn Diagrams**

Differentially expressed genes were acquired from the normalized log2FC data created by DESeq2. Separating by cell line, the over and under-expressed genes for each condition comparison (0RvsControl, 8RvsControl, 8Rvs0R) were categorized as log2FC > 0.5 and log2FC < 0.5, respectively. These were plotted on a histogram in R using ggplot to visualize the number of DEGs present within a cell line at each condition comparison. To visualize both degrees of expression and significance, the same log2FC data was used as input in R to develop a volcano plot using tidy verse’s ggplot2 library. Separate volcano plots were created to compare each condition within each cell line. To visualize the significant batch conserved DEGs within each cell line at a given condition, Venn diagrams were created using tidy verse’s ggplot2 library. Significant DEGs were defined as having a |log2FC| > 0.5 and a p.adj.<0.05. The lists of batches 1 and 2 DEGs for a given cell line and condition comparison were input into ggplot2 to create the Venn diagram. The batch conserved genes were compiled into separate lists. To determine the significant batch and cell line conserved DEGs, the batch conserved lists of genes were separated into UP and DOWN lists, with UP including genes with a log2FC>0.5 and DOWN including genes with a log2FC<0.5. These newly created lists were input into tidyverse’s ggplot2 to make Venn diagrams showing the genes that were similarly expressed in both batches at a given cell line and condition comparison.

**Functional Enrichment Analysis Using GSEA**

Gene Set Enrichment Analysis (GSEA) was run against the ontological gene set collections (C5) defined by the molecular signatures database [50, 51]. GSEA ranks the log2FC data from DESeq2 and designates an enrichment score for each pathway within the gene set collection based on the distance from the middle of this ranked list. For this study, only human collections were used. GSEA normalizes this score to more appropriately compare the different genesets. For this reason, the normalized enrichment score was used in this analysis.

As input for GSEA, an ordered list of l2FC values and corresponding ensemble gene IDs for each batch, cell line, and condition comparison were used. Specific parameters used were “C5” for the geneset to show yield gen ontology gene sets and “Homo Sapiens” as the organism to ensure only human gene sets were queried. GSEA was run using the clusterProfiler library within R. GSEA output included geneset IDs, gene set descriptions, gene set sizes, normalized enrichment scores, p-adjusted values, leading-edge statistics, and lists of the genes associated with each gene set. These results were visualized using the gseaplot2 function and included dot plots, ridge plots, enrichment maps for each batch, cell line, and condition comparison. The dot plots included the ten genesets with the highest normalized enrichment scores and those with the lowest normalized enrichment scores. Dot color denoted p.adjusted value, dot size denoted the number of DEGs within the geneset present in the ordered log2FC list, and placement of dot relative to the y-axis denoted the ratio of differentially expressed genes present concerning the total number of genes within the gene set. The ridgeplots and enrichment maps included the ten genesets with the highest normalized enrichment score for a given batch/cell line/condition.

**Functional Enrichment Analysis using STRING**

STRING analysis was conducted to investigate protein-protein interactions across various batches, cell lines, and condition comparisons. The STRING database aggregates, scores, and integrates publicly available information on protein-protein interactions, leveraging these data sources to generate predictive models of interaction networks [52]. We used our differentially expressed gene (DEG) data obtained from DESeq2 to carry out an enrichment analysis. Specifically, genes with a log2 fold change greater than 0.5 and an adjusted p-value of less than 0.05, along with their Ensembl gene IDs from each batch, cell line, and condition comparison, were input into STRING. The analysis yielded a comprehensive table of gene sets detailing the gene set names, descriptions, enrichment scores, and adjusted p-values for those found to be significantly enriched based on the DEG input. To determine conserved pathways, the tables were compared, and genesets present in both batches and all three cell lines at a given condition were compiled in a table.

**Visualization of Gene Expression Patterns Using Heatmaps**

The heatmap analysis was performed to visualize the expression patterns of genes found in “Heat Acclimation” and “Receptor Ligand Activity” because of heat shock. The log2FC values from the variance stabilized transformed dds object created from DESeq2 were used as input. Thirteen genes from “Receptor Ligand Activity” with conserved gene expression in both batches, three cell lines, and three condition comparisons were visualized. All six genes from “Heat Acclimation” were selected for heatmap analysis. Heatmaps were separately made for each batch to account for the inherent batch effect using the heatmap and BiomaRt libraries [49, 58].

**Cytoscape Generated Network**

Cytoscape's predicted network analysis was performed to determine the interactions of genes based on the acquired expression data. The network analysis uses a list of DEGs and the known interaction networks in the MSigdbr database. We used our list of conserved genes found in “Receptor Ligand Activity” as our list of DEGs. Cytoscape then outputted the predicted interactions of these genes based on available data.
